# NeuRoDev resolves lifelong temporal and cellular variation in human cortical gene expression

**DOI:** 10.64898/2025.12.16.694589

**Authors:** Asia Zonca, Erik Bot, Jose Davila-Velderrain

**Affiliations:** Human Technopole, Viale Rita Levi-Montalcini 1, 20157, Milan, Italy

## Abstract

Understanding how the human brain develops and functions requires direct analysis of human cells. Single-cell atlases open unprecedented opportunities to survey cell physiological molecular states as the brain develops. However, technical challenges limit their potential. We present NeuRoDev, a computational resource with highly curated transcriptomic data and novel analytical tools to investigate neuronal and glial development in the human cortex. NeuRoDev compresses ∼1M single-cell transcriptomes into integrative summary networks of reproducible cell clusters that capture temporal and cellular variation across all stages of human brain development. It provides a reference framework to directly interrogate cellular maturation dynamics, contextualize gene function, and interpret experimental organoid models. We use NeuRoDev to investigate developmental variation in cell physiology, reconstruct genesis and maturation dynamics in neuronal and glial cells, and interpret time-series data from organoid models. NeuRoDev is provided as a freely available software package and as web applications for interactive data analysis.

## INTRODUCTION

The human cortex consists of cells of two main classes: neurons and glia. Neurons are excitable cells that mediate rapid electrical signaling in all nervous systems. In central nervous systems (CNS), neurons form part of functional units together with glial cells that support neuronal function but also sense and process information^1^. Both neurons and glia are highly heterogeneous and plastic in phenotypic state and temporal patterns of appearance and maturation, with evidence of functional conservation and specialization across development^2–4^. Disentangling these processes at cellular and molecular level in a protracted (decades-long) and experimentally inaccessible human brain development has been technically and conceptually challenging^5^. Tissue-resolution profiling studies set the stage by surveying the molecular architecture of the human brain in time and space with high-sampling resolution^6–8^. However, bulk analysis of heterogeneous tissue masks changes in cellular diversity.

Single-cell profiling has addressed this issue and enabled many independent studies to molecularly survey the fetal and adult human cortex at cellular resolution^9–21^. With all these data at hand, many opportunities exist to revisit basic questions in neurobiology and to investigate the cellular and molecular bases of brain disorders in the context of human biology. However, technical and analytical limitations in data access and representation make it difficult to disentangle cellular and temporal changes across decades of cortical development.

Here, we present NeuRoDev, a computational resource that makes use of extensive existing data to resolve temporal and cellular variability during the lifelong process of human cortex development. The motivation behind NeuRoDev is to provide a highly curated data resource and analytical tools to facilitate the study of brain cell physiology in the context of human cortical development. This is achieved by introducing an analytical approach based on integrative networks that connect reproducible transcriptional patterns that compress many single-cell transcriptomes. The resulting networks and summary transcriptomes establish a reference framework to contextualize gene function and iPSC-derived experimental models within human neurodevelopment. We use NeuRoDev to investigate the dynamics of cell physiological processes, the genesis and maturation of neuronal and glial cells; and to interpret data from human experimental models. All analytical tools and data resources are freely available in the R package neuRoDev.

## RESULTS

### Single-cell transcriptomic compendium of human cortical data

To develop the NeuRoDev resource, we first compiled and curated 1,617,236 single-cell transcriptomes from 11 representative studies that collectively survey all stages of cortical development -- from post-conception week 5 to 64 years of age^9–21^ (**Supplementary Table S1, Supplementary Fig. S1a**). We preprocessed the data to obtain 119 individual samples balanced in cell number and labeled with age (n=79), developmental stage (n=13), cell class (n=3), and cell subclass (n=11) (Methods) (**Fig. 1a, Supplementary Fig. S1b**). All developmental stages are represented by at least ∼17k single-cell transcriptomes, with a median of 63,542 cells per stage (**Fig. 1b**). We adopted a staging convention put forward by the BrainSpan consortium based on 13 developmental milestones (https://www.brainspan.org/) (**Supplementary Table S1**). When aggregated by age or stage, samples capture changes in the frequency of cell classes expected from a sequential process of fetal neurogenesis and gliogenesis, with progressive depletion of fetal glia acting as stem cells (radial glia, RG)^4^ and a concomitant increase of neuronal cells (excitatory neurons, ExN) (**Fig. 1c, Supplementary Fig. S1c**). Mature macroglial cells (astrocytes, Ast; oligodendrocyte progenitor cells, OPC; and oligodendrocytes, Oli) increase at late prenatal and early postnatal stages and collectively maintain a high proportion in the mature cortex. Microglial cells are already detected very early in development and persist through life. Few adult samples show depletion of glial cells due to neuronal enrichment in the original study^20^. Expression patterns of glial markers VIM (prenatal samples), GFAP (postnatal samples), and neuronal marker SYT1 (all samples) corroborate these patterns at single-cell level (**Fig. 1d**). Consensus subclass labels across datasets accurately match cell population structure in individual cell atlases, as evidenced by projections on individual 2D cellular landscapes (**Supplementary Fig. S1d,e**). This representative compendium is the major data source used to develop NeuRoDev (Supplementary Data).

**Figure 1.**
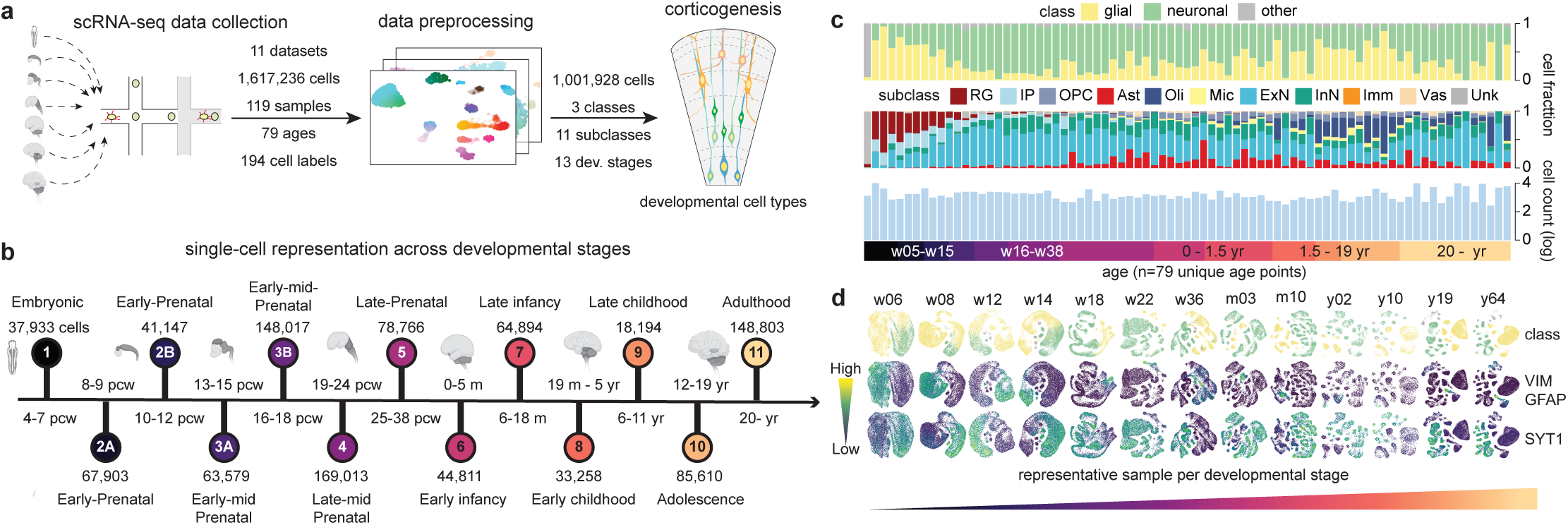
Single-cell transcriptomic data compendium of human developing cortex. **a** Data compilation and preprocessing steps. **b** Representation of the data compendium across 13 developmental stages and corresponding age range. Stages are labeled by numbers (circle) and color-coded. **c** Class and subclass cell proportions and average cell count (log_10_) per individual age data point (columns). Each bar represents samples aggregated according to age (n=79 unique ages). **d** 2D-UMAP representation of the single-cell landscape of one representative sample per developmental stage. Data points represent single cells. Cells are labeled by cell class (top) or by the level of expression of VIM/GFAP (middle) or SYT1 (bottom). VIM was used in prenatal samples to mark glial cells (w) and GFAP in postnatal samples (m-y) to mark astroglia. SYT1 was used in all stages to mark neuronal cells.

### Integrative networks reconstruct cellular and temporal relationships in the cortex

To address the technical limitations implicit in multistudy data integration, NeuRoDev introduces the use of integrative summary networks. The idea is to simplify the analysis and address technical challenges by integrating a reduced number of dominant transcriptional patterns instead of many noisy single-cell profiles. Each sample is compressed into clusters of cells that are represented by their relative expression signatures. These signatures are then integrated across all samples into a single summary network that connects clusters based on their similarity (Methods) (**Supplementary Fig. S2a**). To explore the feasibility of this approach, we first constructed a network that summarizes all neuronal and glial cells in the compendium. The resulting corticogenesis network connects 1,575 high-quality clusters that aggregate in discernible communities of similar cell type and stage and not by study (**Fig. 2a, Supplementary Fig. S2b**). Each cluster summarizes a median of 439 single-cell transcriptomes (19 to 6,141 range).

**Figure 2.**
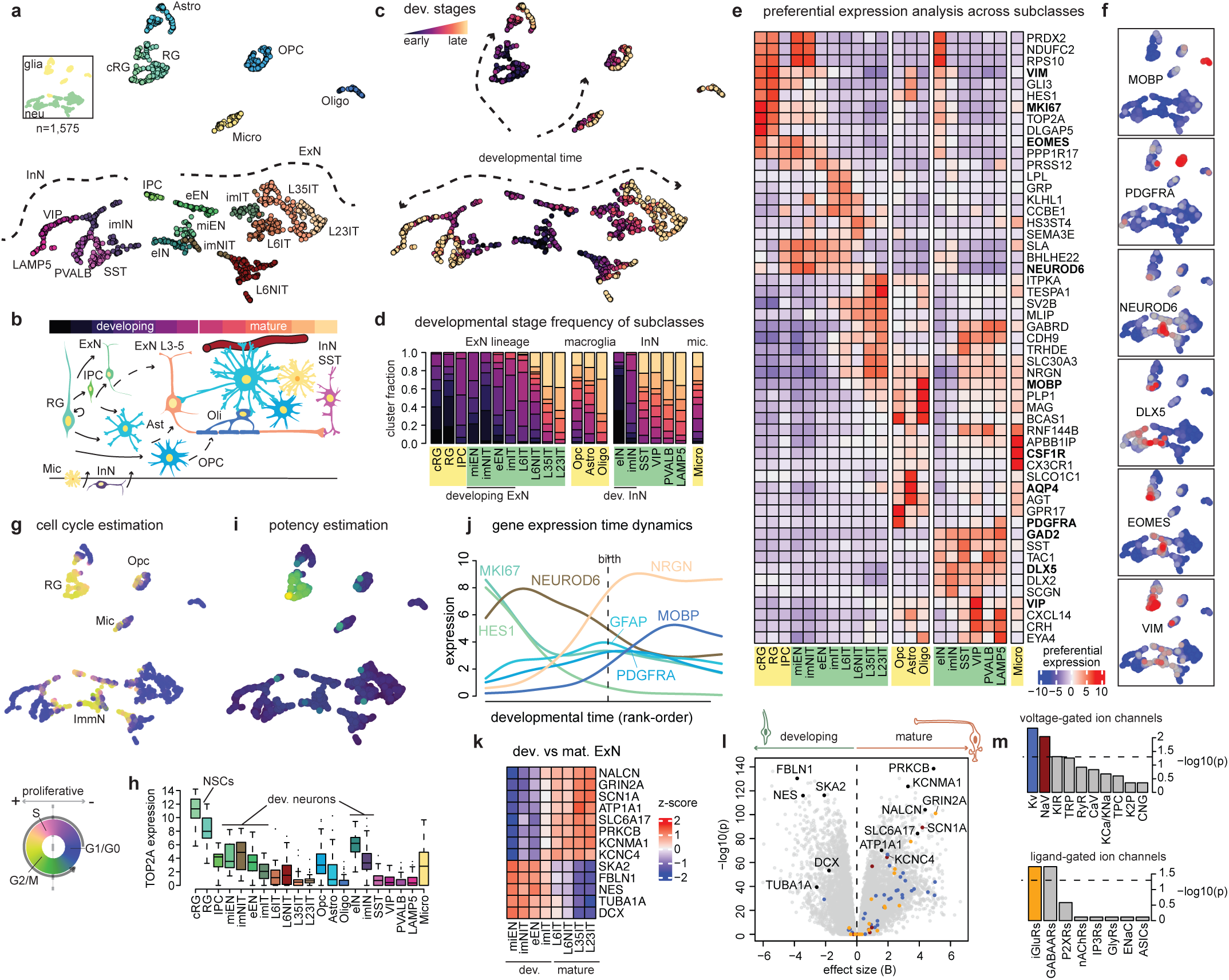
Integrative corticogenesis summary network captures temporal and cellular variability. **a** 2D-UMAP representation of the corticogenesis reference network colored by class (insert) and subclass. Data points represent single clusters (n=1,575). **b** Schematic of main cellular developmental processes. **c** Same as (**a**) colored by developmental stage. Arrows indicate general developmental trends. **d** Developmental stage distribution per subclass. Data shows the fraction of clusters in a developmental stage for each subclass. Subclass labels are color-coded by class (green=neuronal, yellow=glia). Subclasses in each lineage are ordered based on age. **e** Subclass preferential expression profiles of top genes (top 3 per subclass). **f** Expression enrichment values of selected marker genes across the network. Red = high, blue = low. **g** 2D-UMAP representation colored based on cell cycle estimation using tricycle^25^. **h** TOP2A log-normalized expression across subclasses. Data points represent individual clusters: cRG (n=67), RG (n=97), IPC (n=30), miEN (n=42), imNIT (n=36), eEN (n=45), imIT (n=44), L6IT (n=105), L6NIT (n=140), L35IT (n=169), L23IT (n=91), Opc (n=106), Astro (n=103), Oligo (n=58), eIN (n=47), imIN (n=86), SST (n=79), VIP (n=68), PVALB (n=52), LAMP5 (n=33), Micro (n=77). **i)** 2D-UMAP representation colored based on potency scores estimated with CytoTRACE2: brighter colors indicate higher potency. **j** Log-normalized gene expression of selected marker genes across developmental time. Lines are colored by the subclass with highest expression. **k** Relative expression (row z-scaled) of representative genes differentially expressed in mature (mat.) versus developing (dev.) excitatory neurons. **l** Volcano plot comparing expression levels in mature (n=505) versus developing (n=167) excitatory neurons. Data points show differential statistics of individual genes. Voltage-gated potassium (Kv) and sodium (NaV) ion channels (Kv and are shown in blue and bordeaux. Ionotropic glutamate receptor genes (iGluRs) are shown in orange. **m** Over-representation of voltage-gated (top) and ligand-gated (bottom) ion channel families within genes overexpressed in mature vs developing excitatory neurons. Data shows enrichment scores computed with fgsea. Horizontal dashed line indicates a p-value of 0.05. Kv (n=40), NaV (n=9), Kir (n=15), TRP (n=28), RyR (n=3), CaV (n=10), KCa/KNa (n=8), TPC (n=6), K2P (n=15), CNG (n=10), iGluRs (n=18), GABAARs (n=19), P2XRs (n=7), nAChRs (n=16), IP3Rs (n=3), GlyRs (n=4), ENaC (n=4), ASICs (n=3) genes. Box-plots are centered around the median; the interquartile range (IQR) defines the box; whiskers extend to the largest(smallest) value no further than 1.5 × IQR from the end of the box. NSCs = neural stem cells.

Using network structure, cell type correlations, and marker gene expression, we defined 21 higher-resolution subclasses that cover all major groups of glial and neuronal cells (**Fig. 2a,b**, **Supplementary Fig. S2c,d**) (Methods). The upper part of the network recovers developmental and mature glial cells: radial glia (RG), intermediate progenitors (IPC), astrocytes (Ast), oligodendrocyte progenitor cells (OPC), oligodendrocytes (Oligo), and microglia (Mic). The IPC group connects with the lower part of the network, which recovers a bifurcating neuronal structure separating inhibitory (InN, left) and excitatory neurons (ExN, right). The latter group bifurcates further to define intratelencephalic (IT, light brown) and non-intratelencephalic (NIT, dark brown) groups (**Fig. 2a**). Together with glial-neuronal segregation, a second dominant axis of variation across the network is explained by developmental time, as reflected in a middle-out progressive pattern of connection from early to late stages (**Fig. 2c**). Glial and neuronal subclasses show stage preferences consistent with developmental progression. Both developing glial (RG/progenitor cells) and developing neurons (early eN, immature imN, migrating miN) are localized in the middle of the network and enriched in early stages (dark color) (**Fig. 2a-d**). Mature subclasses recover stereotyped temporal patterns of appearance and enrich in postnatal stages (light color). Within ExN, deep-layer neurons (L6NIT, L6IT) are observed earlier, followed by neurons of more superficial layers (L35IT, L23IT). OPC precede Ast and Oligo in stage distribution within macroglia (**Fig. 2d**). Our compendium focuses on primary cortical samples, and thus developing InN cells (eIN, imIN) may represent tangentially migrating cells and/or ventral telencephalon tissue present at the earliest stages of sampling (e.g., w05-w08). Mature InN neurons recover major cardinal classes observed in the adult cortex (PV, VIP, SST, LAMP5)^22^. The non-ectodermal microglia show a broad developmental distribution. Mic clusters are observed already at the first stages and continue throughout development, consistent with the early establishment of microglia in the CNS^23^.

To directly compare expression changes across subclasses and stages, we generated a dataset of 1,575 pseudobulk transcriptomes that complement the network (Methods). These profiles capture cell physiological states and facilitate direct comparison with external data sources. Top preferentially expressed genes in each subclass identify markers routinely used for cell type identification (**Fig. 2e**). Expression of these markers overlaps with subclass distributions across the network (e.g., VIM in RGs, EOMES in IPC, NEUROD6 and DLX5 in dorsal and ventral developing neurons, PDGFRA in OPC, and MOBP in Oligo) (**Fig. 2f, Supplementary Fig. S2d**). Developmental stage changes at pseudobulk level correlate with temporal expression patterns of bulk cortical tissue at expected times, supporting NeuRoDev’s age-based staging (**Supplementary Fig. S2e**). Prenatal ExN lineage subclasses correlate with microdissected transcriptomes of matching cortical germinal zones^6^ (**Supplementary Fig. S2f**). RGs and progenitors correlate with (sub)ventricular zones (VZ, iSZ, oSZ), developing neurons with subventricular zones (iSZ, oSZ) and cortical plate (iCP, oCP), and mature neurons with upper lamina. Profiles of mature ExN subclasses match transcriptomes of corresponding cortical layers in spatial transcriptomics data^24^ (**Supplementary Fig. S2g**). Estimation of cell cycle position from pseudobulk profiles with tricycle^25^ predicts proliferative states in RG cells, consistent with their neuronal stem cell function in the fetal brain^4,26^, and in subpopulations of OPC and microglial cells, the most proliferative mature glia in CNS^23^ (**Fig. 2g**). The transcriptome of early immature cells adopting a neuronal fate (e.g., eIN, imNIT) also predicts proliferative states, which gradually become postmitotic along maturation trajectories in the network. These predictions are consistent with pseudobulk expression patterns of topoisomerase TOP2A, a key controller of DNA topology during replication^25^, and with the specificity and temporal dynamics of the proliferation marker MKI67 (**Fig. 2h, j**). Glial proliferative clusters overlap with pseudobulk estimates of potency obtained with CytoTRACE2^27^ (**Fig. 2i**). Transcriptional states of neuronal cells are not predictive of potency, irrespective of maturation, consistent with a restricted fate at early developmental stages despite ongoing cell cycling activity in the most immature. The cell type specificity and temporal expression of representative markers are consistent with sequential progression of neurogenesis and gliogenesis, with progressive reduction of proliferative (MKI67) and neuronal stem cell (HES1) markers, and sequential peaking of proneuronal (NEUROD6), astrocyte (GFAP), oligodendrocyte precursor (PDGFRA), neuronal (NRGN), and mature oligodendrocyte (MOBP) markers (**Fig. 2j**).

Differential expression analysis of mature versus developing ExNs revealed a molecular phenotype suggestive of functional immaturity in the latter (**Fig. 2k-m**). Mature neurons overexpress molecular complements required for electrical signaling, neurotransmission, and calcium-mediated signal transduction, including pore-forming alpha subunits of the voltage-gated sodium channels (NaV, SCN1A), subunits of voltage-gated (Kv, KCNC4) and calcium-activating (KCa, KCNMA1) potassium channels, the sodium leak channel (NALCN), the glutamine transporter SLC6A17, and ionotropic glutamate receptors (iGluRs, GRIN2A) (**Fig. 2k-m**). In contrast, developing neurons overexpress the immature markers Doublecortin (DCX) and Nestin (NES), together with genes involved in neuronal migration (TUBA1A, FBLN1) and the onset of mitosis (e.g., SKA2). Lastly, top preferentially expressed biological processes estimated directly from pseudobulk across subclasses (Methods) support these observations and broadly recover cellular physiological expectations. For example, profiles show preferential mitotic DNA replication in RGs, neuronal migration and collateral sprouting in developing neurons, neuronal ion channel clustering and synaptic vesicle docking in mature neurons, GABA transport in InN, GPCR-mediated glutamate signaling in ExN, axon ensheathment in oligodendroglia, and glucan biosynthesis in Ast (**Supplementary Fig. S2h**). Taken together, these data support the reliability of integrative summary networks and profiles as interpretable frameworks to investigate cellular and temporal variability in the human cortex.

### NeuRoDev traces temporal and cellular variation and recovers cell physiology

NeuRoDev introduces a tool that allows direct visualization of cellular and temporal expression variability across cortex development. We call this tool eTrace (expression Trace) analysis. An eTrace defines a reference space that traces expression-derived molecular phenotypes (y-axis) in time (x-axis), while tracking developmental stage and cell identity with colors (**Fig. 3a**). Expression of cell type marker genes in this space identifies clusters corresponding to cells of the expected type (**Fig. 3b**). Because eTraces compress both cell and time (age) information, the analysis evidences whether the expression phenotype is time-restricted -- i.e., observed only at certain stages. It also allows direct estimation of developmental trends of gene expression (red lines) (**Fig. 3b**). Time restriction can result simply because the cell type is only present at certain stages (e.g., PLP1-enriched mature oligodendrocytes), but it can also relate to maturity or developmental variation within cell types (e.g., NRGN is only enriched in mature neurons, while InN GAD2 enrichment is time-invariant) (**Supplementary Fig. S3a)**. Other possible patterns involving nontrivial time-cell type relationships are those associated with lineage and/or conservation of function. For example, astroglia includes both developing and mature cell types that cover all developmental stages and involve developmentally conserved functions of neuronal support as well as cell type and temporal specializations^4,28^. eTrace analysis can intuitively capture this behavior. For example, while the enrichment of the canonical RG marker VIM is developmentally restricted in RG, the markers PAX6 and GLI3 are conserved in astroglia throughout development, with enriched expression in both RGs and astrocytes (**Fig. 3b, Supplementary Fig. S3a)**.

**Figure 3.**
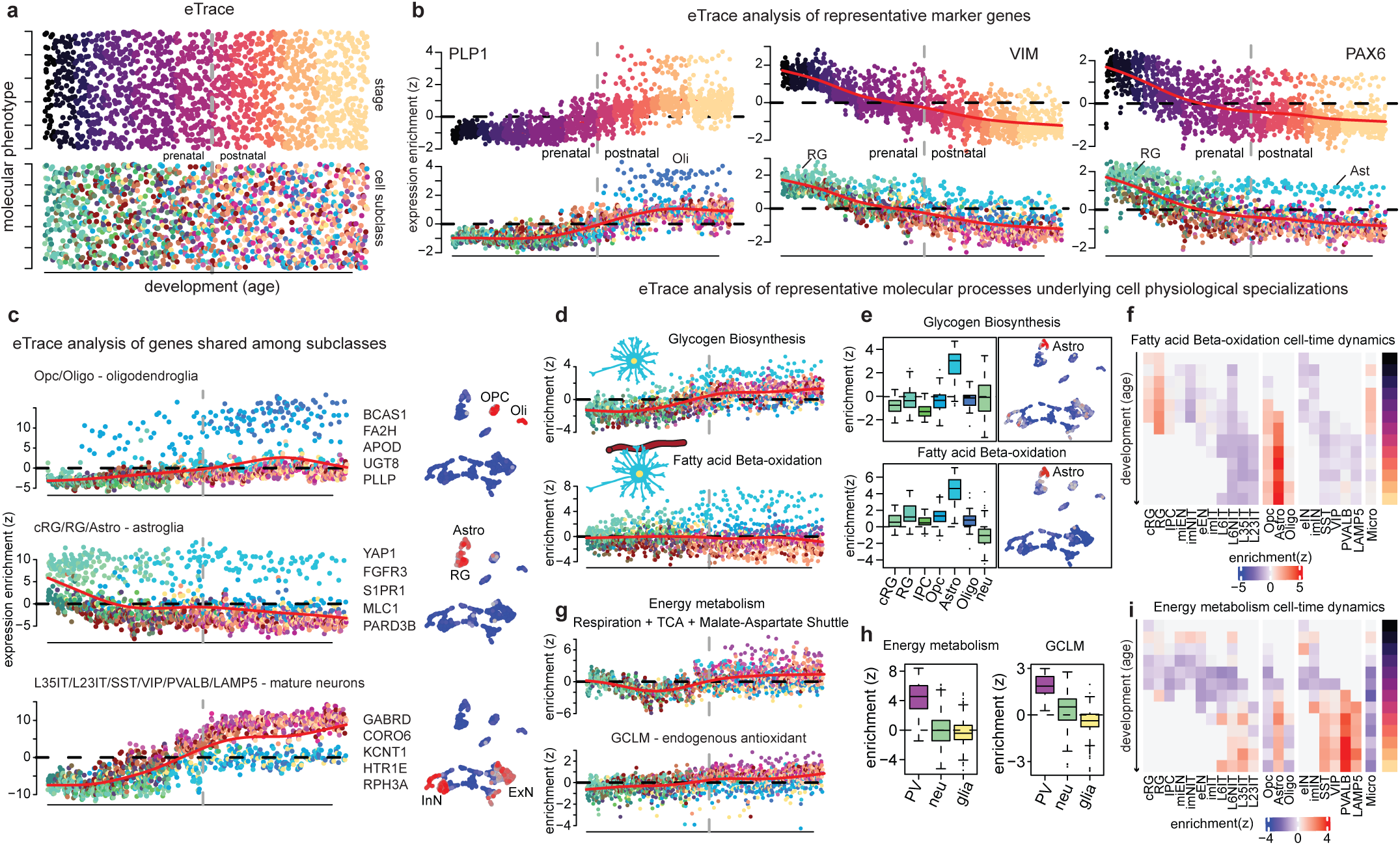
eTrace analysis to investigate temporal and cellular expression patterns. **a** eTrace schematic representation. Points represent single clusters colored by stage (top) and subclass (bottom) and ordered incrementally by age (x-axis) and mapping values of a transcriptionally derived molecular phenotype (y-axis). The vertical grey dashed line represents birth. **b** eTrace analysis of selected marker genes: PLP1 (Oli), VIM (RG), PAX6 (RG, Ast) (n=1,575 clusters). Red lines represent temporal trends. **c** eTrace analysis (left) and expression enrichment network projection (right; red = high, blue = low) of genes preferentially expressed in the oligodendroglia lineage (top), the astroglia lineage (middle), or mature neurons (bottom). Five representative genes are shown for each category (middle). **d** eTrace analysis of genes involved in glycogen biosynthesis (top) and fatty acid beta-oxidation (bottom). **e** Distribution of expression enrichment values across subclasses (left) and projected across the network (right) (red = high, blue = low). Progenitors (cRG, RG, IPC), astrocytes (Astro), oligodendrocytes (Oligo), neurons (neu). **f** Expression enrichment values of genes involved in fatty acid beta-oxidation across subclasses (columns) and developmental stages (rows). **g** Same as (**d)** for genes involved in energy metabolism (top) and for the GCLM gene (bottom). **h** Distribution expression enrichment values of genes involved in energy metabolism (left) and the GCLM gene (right). Parvalbumin neurons (PV) (violet), other neurons (green), glia (yellow). **i** Same as (**f**) for genes involved in energy metabolism. Box-plots are centered around the median; the interquartile range (IQR) defines the box; whiskers extend to the largest(smallest) value no further than 1.5 × IQR from the end of the box.

To further explore the ability of eTrace analysis to capture time-cell relationships, we systematically identified 2,469 genes that are preferentially expressed by one (exclusive) or several (shared) cell subclasses (Methods) (**Supplementary Table S2**). Consistent with the high heterogeneity and specialization of glial cells^3^, non-ectodermal microglia have the largest number of exclusive genes, followed by mature glia (Oligo, Ast, OPC) and cRG. Individual neuron subclasses, in turn, have low numbers of exclusive genes because they tend to share “neuronal” genes across several or most subclasses, but not with glia (**Supplementary Fig. S3b)**. Consistent with lineage relationships, RG subclasses (cRG and RG) share the most genes, followed by IT neurons (L35IT, L23IT), oligodendroglia (OPC, Oligo), subsets of mature neurons (e.g., InN subclasses), astroglia (cRG, RG, Ast), progenitors and developing ExN neurons, and macroglia (OPC, Astro, Oligo) (**Supplementary Fig. S3b**). To directly visualize the behavior of these genes on eTraces, we extended the analysis beyond individual genes to include gene set enrichment traces, relative to random behavior (Methods). eTraces of subclass-exclusive genes separate the involved cell types with high accuracy, and their enrichment patterns highlight network neighborhoods that match expected subclasses (**Supplementary Fig. S3c**). eTraces of gene groups shared among subclasses intuitively recover time-cell and lineage relationships segregating macroglial, oligodendroglial, astroglial, and neuronal signatures (**Fig. 3c, Supplementary Fig. S3d**). eTrace analysis thus provides an easy way to map enrichment patterns of individual genes and gene groups across cell subclasses and developmental stages.

We next sought to investigate whether eTrace analysis easily captures molecular processes associated with known cell physiological specializations. Despite its high energy requirements, the brain has almost no energy reserves, except for glycogen storage in astrocytes^29,30^. eTrace analysis of glycogen biosynthetic pathways recovered this specialization, with a pronounced and specific enrichment in Ast (**Fig. 3d**). Similarly, the documented glycolytic preference of astrocytes^30^ is readily recovered by eTrace analysis of glycolytic pathway genes, although with lower specificity and with slight enrichment in RGs (**Supplementary Fig. S3e**). To generalize these observations and explore potentially novel associations, we identified biological processes, molecular functions, and cellular components that are preferentially expressed among subclasses (Methods) (**Supplementary Table S2**). Preferential processes generally match known cell physiology. For example, mitotic processes are enriched in RGs and IPCs; sprouting, migration, and translation in developing neurons; neurotransmitter receptors and voltage-gated ion channels in mature neurons; and metabolic and myelinating pathways in mature glia (**Supplementary Fig. S3f**). Together with these expectations, we also identified additional notable associations in glia and neurons. Consistent with the functional role of astrocytic fatty acid metabolism *in vivo*^31^, multiple pathways involved in beta oxidation are specifically enriched in Ast (**Supplementary Fig. S2g**). eTrace analysis of genes involved in beta oxidation, including transporters (ABCD2/3/4) and enzymes (ACAD/L/VL/M), confirmed this association and identified significant enrichment in mature astroglial cells (**Fig. 3d-f**). Lastly, we found a neuronal association between fast-spiking PV-expressing interneurons (PVALB InNs) and energy metabolism. The fast-spiking properties of these neurons impose high metabolic demand and increased mitochondrial density^32^. In agreement with this physiological demand, the PVALB subclass shows enriched expression of several processes implicated in energy metabolism, including *Respiratory Chain*, *TCA cycle*, *ADP Transport*, and the *Malate−Aspartate Shuttle*. eTrace behavior of these genes highlights PVALB clusters with higher enrichment over other neuronal or glial subclasses (**Fig. 3g-i**). This is accompanied by a similar eTrace behavior of GCLM, a rate-limiting enzyme of the endogenous antioxidant and redox regulator glutathione (GSH), and one possible contributor in tolerating the oxidative stress imposed by high metabolic demand^33^ (**Fig. 3g)**. Taken together, these data support eTrace analysis as a useful framework to directly study temporal and cellular variability across cortical development. It is able to recover well-established physiological roles of different cell types and potentially novel gene and pathway associations.

### NeuRoDev integrates a neuronal genesis and maturation network

To resolve variation within developing neuronal and glial cells with higher resolution, NeuRoDev includes lineage-specific integrative networks. The rationale for developing these networks is the potential to uncover subtle differences within progenitors, neurons, and glia that are not apparent when analysing cells together. We generated a neuronal development network (neurogenesis) that connects 1,248 high-quality clusters identified *de novo* per sample considering all RG, progenitor, and ExN cells in the reference compendium. We complemented the network with 1,248 pseudobulk transcriptomes used to compare expression levels directly and identify 1,265 genes preferentially expressed by one or several subclasses and corresponding gene ontology (GO) processes (**Supplementary Fig. S4a,b, Supplementary Table S2**) (Methods). The network recovered 25 higher-resolution subclasses annotated based on marker genes, stage/cell relationships, and transcriptional patterns (**Fig. 4a, Supplementary Fig. S4c-e**). The lineage-specific analysis identified additional subclasses of RG/progenitor cells interpreted as ventricular (vRG, VIM+), cycling (cRG, TOP2A+), or outer (oRG, TNC+); an early subclass of IPCs (eIPC, NEUROG1+); and a late subclass of RGs (lRG) present at late prenatal stages and expressing mixed markers of RG/progenitor subclasses. Neurons include 7 developing subclasses interpreted as migratory (miNIT), immature (imNIT, imIT1/2), newborn (nExN), or early (eL6B, eL6CT); and 12 mature ExN subclasses annotated with putative layer (L2/6) and projection pattern (extratelencephalic ET, intratelencephalic IT, corticothalamic CT, near-projecting NP). Complete label descriptions are reported in **Supplementary Table S3**. Developing neurons preferentially express semaphorins (SEMA3C, SEMA3E), chemorepulsive proteins (DRAXIN), actin-binding proteins (ACTN2), and synaptotagmins (SYT6); consistent with maturation processes involving neuronal and axon guidance, acting remodeling, and Ca^2+^-dependent sensing (**Supplementary Fig. S4c**).

**Figure 4.**
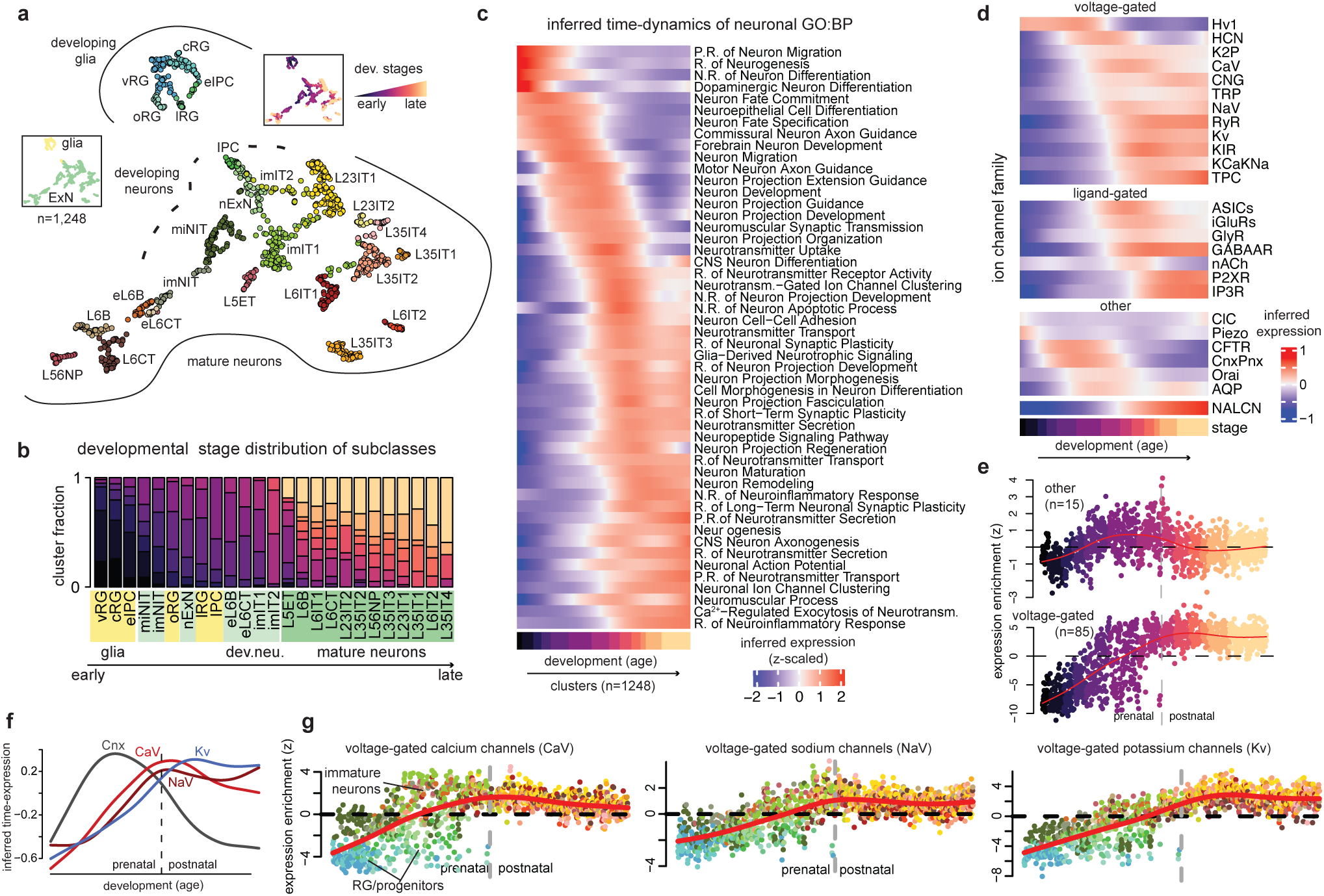
Neuronal genesis and maturation network. **a** 2D-UMAP representation of the neurogenesis reference network colored by subclass, class (left insert), and developmental stage (right insert). Data points represent individual clusters (n=1,248). **b** Developmental stage distribution per subclass. Data shows the fraction of clusters in a developmental stage for each subclass. Subclass labels are color-coded by group (green=mature neurons, light green=developing neurons, yellow=glia). Subclasses are ordered based on age. **c** Temporal expression trends of neuronal Gene Ontology biological processes across clusters. Clusters are ordered by age. **d** Temporal expression trends of ion channel families across age-ordered clusters. **e** eTrace analysis of voltage-gated ion channels (bottom) and channels not gated by voltage or ligands (top) colored by developmental stage. **f** Inferred expression trends of voltage-gated sodium (NaV), potassium (Kv), and calcium (CaV) ion channels, and connexins (Cnx) across development. **g** eTrace analysis of voltage-gated calcium (left), sodium (center), and potassium (right) channels, colored by subclass. R. = regulation, P. = positive, N. = negative, Neurotransm. = neurotransmitter.

To systematically investigate the consistency of our data and annotations with temporal changes in known maturation processes, we compiled neuronal GO:BP processes (n=79 terms, 2,702 genes) and compared their expression across subclasses and time (Methods) (**Fig. 4c**). Transient expression waves mapped neuronal differentiation, fate commitment, and specification to early developmental stages and RG/progenitor and immature neuron subclasses. Neuronal migration and axon/projection extension, guidance, and organization mapped to late prenatal and early postnatal stages and developing neuron subclasses. A sustained expression wave mapped neurotransmitter transport, secretion, and signaling; neuron maturation; neuronal ion channel clustering and action potential to postnatal stages and mature neurons, with a parallel increasing pattern of neuroinflammatory responses (**Fig. 4c**). The core processes of mitosis, DNA repair, and translation are enriched early, followed by axon guidance and chemotaxis, and finally by synaptic transmission. These processes match the sequential enrichment of nuclear, ribosomal, and cytoskeletal cellular components, followed by axons, dendrites, and synapses (**Supplementary Fig. S4f**). The waves observed in our integrative human data are consistent with enrichment patterns of broad biological processes and cellular structures previously shown to mark the developmental progression of maturing neurons labelled *in vivo* in mice^34^.

To further explore the extent to which our integrative data captures aspects of the physiological maturation of human developing neurons, we next analysed the temporal expression patterns of different kinds of ion channels (Methods). The diversity of ion channels expressed by a neuron largely determines its potential to encode and transmit complex signals^35^, and their developmental activity and regulation impact neuronal differentiation^36,37^. Most ion channel families (26 out of 30) express at least one subunit at some timepoint in our neurogenesis network (log_2_(CPM) > 2) (**Fig. 4d**, **Supplementary Fig. S4g**). Calcium-Dependent Chloride Channel (CaCC), 5-HT3 receptors, epithelial sodium channels (ENaC), and Zinc-activated channels (ZAC) were not detected in our data. Voltage-gated ion channels (VGICs) provide the molecular basis of neuronal excitability^38^ and largely determine the intrinsic encoding properties of mature neurons^39,40^. VGICs also contribute to the spontaneous electrical activity of developing excitable cells^41^. Members of all VGIC families are expressed in our network, with increasing overall expression as neurons mature (**Fig. 4d**). Unlike most VGICs, the proton-selective channel Hv1 (HVCN1) is expressed early during neurogenesis, with higher expression in RGs and immature neurons (**Supplementary Fig. S4g**). Other early enriched ion channels are not gated by voltage or ligands and include mechanosensors (Piezo) and gap-junction channels (CnxPnx), in agreement with the role of cellular adhesion and gap junctional communication in cytoskeletal organization and radial migration^42^ (**Fig. 4d, Supplementary Fig. S4g**). eTrace analysis of neuronal development further suggests prenatal enrichment of ion channels not gated by voltage/ligands (other), at a time when cytoarchitectural structure is being formed and immature neurons undergo migration. Channels gated by ligands or voltage begin to express prenatally but enrich throughout postnatal development, when mature neurons are chemically and electrically coupled in circuits (**Fig. 4e, Supplementary Fig. S4h**). Within VGICs, our developmental transcriptomic data suggest sequential aggregate expression of calcium (CaV), sodium (NaV), and potassium (Kv) channels as neurons mature (**Fig. 4f,g, Supplementary Fig. S4g,i**). Developing neurons at mid and late prenatal stages show enrichment of CaV channels (**Fig. 4g**). Both CaV and NaV channels are expressed early in neuronal differentiation, while Kv expression peaks postnatally (**Fig. 4f**). These global patterns are explained at gene level by differences in expression specificity of subunit encoding genes. Most CaV subunits show constant expression starting early and throughout development. In contrast, genes encoding NaV and Kv subunits show higher developmental variability, with changes in expression levels in some subunits and time-restricted expression in others (**Supplementary Fig. S4i**). Differential developmental specificity among subunits is observed within multiple families (e.g., KIR, HCN, K2P), and cortical developing neurons seem to progressively acquire a diverse ion channel complement (**Supplementary Fig. S4g**). These observations are consistent with a dynamic development of different ion currents as observed functionally in experimental models^36,43^ and suggest our data could be useful in investigating transcriptional changes underlying the *in vivo* electrophysiological maturation of human cortical neurons.

### NeuRoDev integrates a glial development network

To facilitate higher-resolution analysis of glial cells, we generated a glial development (gliogenesis) network connecting 785 clusters identified by independently analyzing all RG, progenitor, and mature glial cells of the compendium (Methods) (**Fig. 5a**). The network is complemented by 785 pseudobulk transcriptomes used to directly compare gene expression and identify 1,558 genes preferentially expressed by one or several glial subclasses and corresponding GO processes (**Supplementary Fig. S5a,b; Supplementary Table S2**). The exclusion of neurons enabled the identification of 18 higher-resolution glial subclasses, including 9 groups of progenitors and 9 of macroglial cells distinguished by developmental distribution and expression profiles. Within progenitors, the network recovered 6 populations of RG cells interpreted based on expression signatures as early (eRG), ventricular (vRG), S phase cycling (sRG), G2M phase cycling (g2mRG), outer (oRG), and truncated (tRG); and 3 groups corresponding to intermediate (IPC), multipotent (MPC), and neuron-primed (NPC) progenitors (**Supplementary Fig. S5c,d**). The NPC group is more frequent at early stages, correlates with signatures of neuronal subclasses, and has the largest number of uniquely preferential genes, consistent with a pre-neuronal state diverging from glia. IPC and MPC are frequent at late prenatal stages, correlate with signatures of both glial and neuronal subclasses, express markers of oligodendrocyte lineage cells (e.g. OLIG2), and have only few or none uniquely preferential genes (n=5 MPC; n=0 IPC) (**Fig. 5b, Supplementary Fig. S5b,d,e**). IPC instead share preferential genes with OPCs, including CSPG4, GALR1, SULF2, CHST15, GSG1L, and CDH13, suggesting an intermediary gliogenic state (**Supplementary Table S2**). Thus, our network likely includes subpopulations of neurogenic, gliogenic, and multipotent RGs^44^.

**Figure 5.**
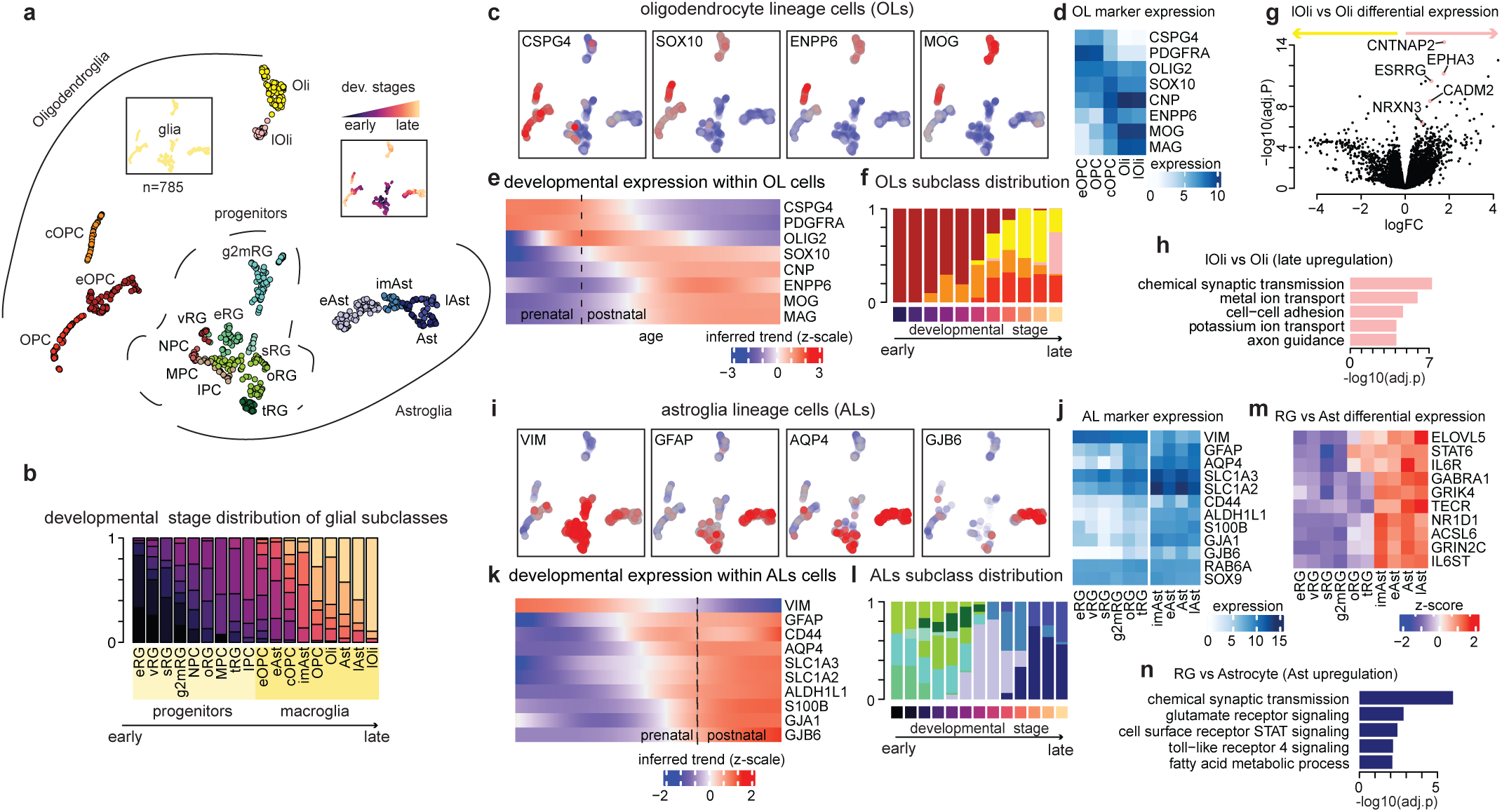
Glial development network. **a** 2D-UMAP representation of the gliogenesis reference network colored by subclass, class (left insert), and developmental stage (right insert). Data points represent single clusters (n=785). **b** Developmental stage distribution per subclass. Data shows the fraction of clusters in a developmental stage for each subclass. Subclass labels are color-coded by developmental class (light yellow=progenitors, yellow=macroglia). Subclasses are ordered by age. **c** Projection of marker gene expression enrichment on the gliogenesis network (red = high, blue = low). **d** Expression of OL markers across OL subclasses. **e** Temporal expression trends of OL markers across OL clusters ordered by age. **f** OL subclass distribution by developmental stage. Data shows the fraction of clusters in a subclass for each developmental stage. **g** Difference in expression levels in lOli versus Oli subclasses. Data points show differential statistics of individual genes. **h** Representative Gene Ontology Biological Processes enriched in lOli compared to Oli. Data shows - log10 of adjusted p-values calculated with fgsea. **i** Same as (**c**) for AL markers. **j** Same as (**d**) for AL markers and subclasses. **k** Same as (**e**) for AL markers. **l** Same as (**f**) for AL subclasses. **m** Relative expression values (row z-scaled) of selected genes differentially expressed in Ast versus RG. **n** Same as (**h**) for Ast versus RG. OL = oligodendroglia lineage; AL = astroglia lineage.

The two principal macroglial cells segregate in two main lineages (**Fig. 5a**). Oligodendrocyte lineage cells (OLs) localize in the upper part of the network and include several OPC and oligodendrocyte states. We interpreted these states based on developmental distribution and marker genes as early, committed, or late (eOPC, cOPC, lOli). In the opposite end (lower-right), astrocytes include early, immature, and late states (eAst, imAst, lAst). OPCs coexpress CSPG4 (NG2), PDGFRA and OL markers like OLIG2 and SOX10 (**Fig. 5c,d**). Temporal expression patterns across the pseudobulk profiles of developing OLs recover known OPC differentiation signatures, with initial expression of OPC and OL markers, followed by differentiation state markers (e.g., CNP, ENPP6), and with myelin sheath proteins of mature oligodendrocytes expressing last (MOG, MAG)^2^ (**Fig. 5e**). These expression patterns reflect developmental changes in cellular composition from an early dominant population of OPC to one dominated by Oli and passing through cOPC cells at intermediate stages (**Fig. 5f**). cOPC show enriched expression of the G protein-coupled receptor GPR17 and the Brain Enriched Myelin Associated Protein 1 (BCAS1), both previously identified in “differentiation committed OPCs”^45,46^, together with lower levels of myelin protein encoding genes (e.g, MOBP, MOG, PLP1) than oligodendrocytes (**Supplementary Fig. S5d**). Oligodendrocytes (Oli and lOli) have the largest number of shared signature genes (**Supplementary Fig. S5b)**, suggesting a highly distinct transcriptome and explaining the extreme separation of these cells in the network. Among signature genes both Oli and lOli preferentially express the connexins Cx32 (GJB1) and Cx47 (GJC2), and the mechanosensor PIEZO2 (**Supplementary Fig. S5g)**. The lOli population additionally expresses higher levels of the cyclin dependent kinase inhibitor CDKN2A. When comparing the transcriptomes of lOli versus Oli cells, the former showed increased expression of genes involved in cell-cell adhesion, chemical synaptic transmission, and ion transport, including CNTNAP2, EPHA3, CADM2, and NRXN3; a profile suggestive of a subset of mature OL cells more readily engaged in functional glial-neuronal interactions (**Fig. 5g,h**).

Astroglia is a diverse class of cells including several cell types^47^. Of these, our network includes RG cells and astrocytes, and for consistency we refer to them collectively as astrocyte lineage cells (ALs) (**Fig. 5i**). The ALs in the network express common astrocyte markers (e.g., VIM, GFAP, AQP4, GJB6/Cx30) with different degrees of specificity and patterned temporal dynamics (**Fig. 5i-k**). All ALs express intermediate filament proteins, with VIM expressed at higher levels at early stages in RG cells, and GFAP in late prenatal RGs and postnatal astrocytes. The aquaporin AQP4 is expressed first at late prenatal stages and maintained throughout postnatal stages and across astrocyte states. The excitatory amino acid transporters SLC1A2 and SLC1A3 are expressed in both RG and astrocytes but increase in expression across development. Both RGs and astrocytes express specific connexins, with both GJA1 (Cx43) and GJB6 (Cx30) expressed more highly in astrocytes and the latter also more specifically (**Fig. 5j,k**). Marker expression and signature gene sharing patterns are consistent with cell population changes across developmental stages (**Fig. 5l, Supplementary Fig. S5b**). RG cells are observed only prenatally in the human developing cortex and subgroup combinations share many preferential genes (e.g., proliferation marker MKI67, centrosome protein genes CENPM, CENPH, CENPU, CENPW, and Rho GTPase activating genes ARHGAP11A). oRG and tRG are observed at late prenatal stages, overlapping with the earliest astrocytes (eAst). Most Ast subclasses are observed postnatally and share signature genes (e.g., ALDH1A1, GJA1, APOE, GNA14). When comparing the transcriptomes of all noncycling RG subclasses versus all Ast subclasses, the overexpression in the latter of genes involved in synaptic transmission (GABRA1), glutamate receptors (GRIK4, GRIN2C), immune-related signaling (STAT, toll-like receptor; STAT6, IL6R), and fatty acid metabolism (ELOVL5, TECR) seems the most prominent difference. Thus, in addition to conserved neuronal support functions in both RG and astrocytes, the differential transcriptome suggests signatures of defense and neurotransmitter (e..g, glutamate, GABA) and systemic homeostasis (e.g., chemosensing)^47^ differentiates the latter. Although we observe expression of astrocyte makers across all subclasses, there are significant differences in expression levels in some cases. The early astrocytes (eAst) express higher levels of filament proteins VIM and NES, genes involved in fatty acid metabolism (FABP7, TSPO), and proliferation markers (MKI67, TOP2A), suggesting an immature, possibly transitory state. The imAst and Ast states express higher levels of neurotransmitter transport and metabolism genes (SLC1A2/3, MAOB) and neurotrophic and Ca^2+^ signaling (NTRK2, NFATC1/2), while the late state (lAst) expresses higher levels of some of the markers commonly associated with reactive astrocytes^48,49^ (e.g., GFAP, CD44, MT2A, MT1E, HSPB1, CRYAB), suggesting some clusters of mature astrocytes possibly recover reproducible reactive states across datasets (**Supplementary Fig. S5h)**. Taken together, our data suggests that the integrative glial development network captures molecular aspects of the high heterogeneity^1,3,4,28^ and developmental plasticity^2^ of human cortical glial cells.

### NeuRoDev contextualizes iPSC-derived human experimental models

Interpreting transcriptomic data from experimental models is challenging and time-consuming, often relying on having access to multiple external reference datasets. To facilitate this process, NeuRoDev enables mapping single-cell or bulk RNA-seq data onto the reference networks, thus providing a toolkit for annotation and biological interpretation of query data via knowledge transfer (**Supplementary Fig. S6a,b**). To illustrate NeuRoDev mapping strategy, we reanalyzed published bulk^50^ and single-cell^51^ RNA-seq data of iPSC-derived human organoids. The bulk dataset includes cortical organoids sampled at 11 different time points ranging from day 75 (d75) to day 600 (d600). We averaged samples from the same time point and mapped them onto the corticogenesis network (**Fig. 6a**) (Methods). To estimate how well each organoid sample aligns with the reference clusters in the network, we assigned a mapping confidence score to each sample (**Supplementary Table S4**) (Methods). The mapping confidence decreased over time (day 25 confidence=0.75; day 600 confidence=0.65), which may indicate that organoids developed for a longer time acquire transcriptional signatures less comparable to those observed *in vivo*. From the 2D-UMAP representation of the corticogenesis network, it is apparent that organoids become increasingly similar to developing neurons as they mature (**Fig. 6a**). However, this trend reverts at later time points (e.g., d200 onwards). To further explore this observation, we interpret each organoid sample in terms of cellular subclasses and developmental stages by transferring knowledge from the network neighborhood and defining annotation scores indicating how similar each sample is to the reference annotations (**Fig. 6b**) (Methods). The subclass distribution shows increasing neuronal similarity from day 25 (d25) to day 200 (d200), followed by a decrease in later time points. The stage distribution further highlights a decreasing similarity to mature cell states after d200, and conversely an increasing similarity to early developmental stages (dark colors) (**Fig. 6b**). Given the proximity between radial glia and astrocyte clusters in the network, and their astroglial relationship, we further analysed organoid samples from late time points to assess which cell type they resemble the most. Similarity to astrocytes would suggest the *in vitro* generation at later time points of cells resembling mature glia, thus recapitulating the sequential neuro-glia genesis observed *in vivo* in the developing cortex. In apparent agreement with this expectation, the expression level of genes specific to astrocytes progressively increases in organoids up to d400, where the expression peaks. However, it subsequently decreases at d500 and d600. This trend is mirrored by the expression of cRG-specific genes, which progressively decrease in expression in the organoids up to d400, and then increase as astrocytic genes decrease (**Fig. 6c**). This analysis thus suggests that later-time organoids acquire a radial glia-like transcriptional signature and not an astrocytic one. This interpretation is also consistent with the transcriptional similarity (average correlation values) between organoids at d600 and the reference subclasses, which shows higher similarity to progenitor and immature cell types than to astrocytes or other mature glial cells (**Fig. 6d**). To investigate how genes preferentially expressed in organoids at d600 normally express during cortical development, we then applied eTrace analysis. The eTrace visualization of these d600 organoid genes confirms high early expression in radial glia clusters, but also a high and time-invariant enrichment in microglia clusters, possibly indicative of cellular inflammation (**Fig. 6e**).

**Figure 6.**
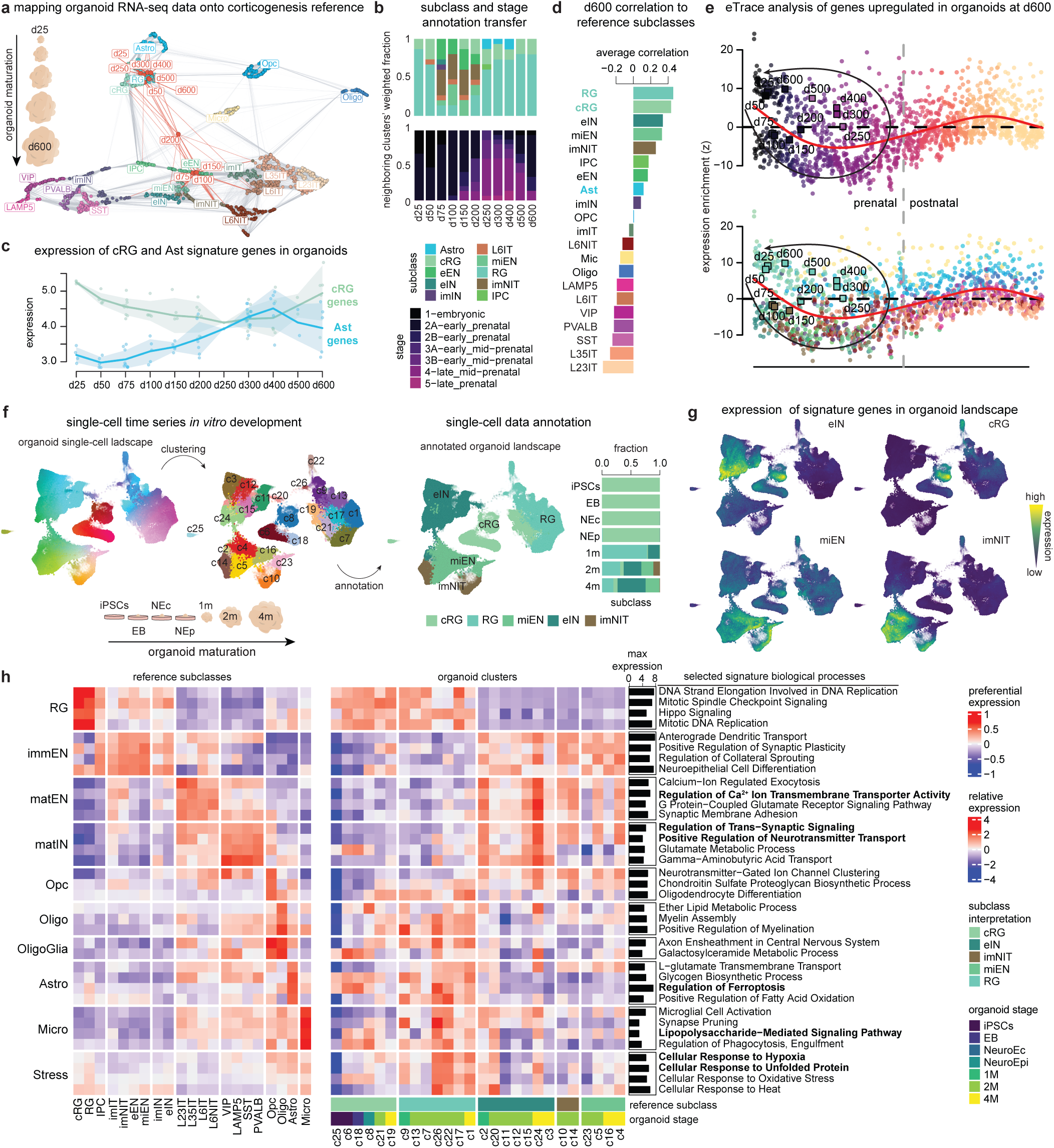
Mapping and interpretation of query organoid data. **a** 2D-UMAP representation of the corticogenesis reference network colored by subclass. Red dots represent query bulk RNA-seq cortical organoid samples^50^ mapped to the network. Labels indicate the day of differentiation of the organoid. Red edges connect query samples to top 15 nearest neighbor reference clusters based on correlation values. Point coordinates are estimated based on similarity to reference clusters. **b** Subclass (top) and stage (bottom) annotation scores (*Methods*) for each query sample. **c** Expression levels in organoid samples of known preferentially expressed genes of cRG (green) and Ast (light blue). Lines represent the mean expression across replicates of the same time point; shaded area represents the mean ± standard deviation. Data points represent single replicates. **d** Transcriptional similarity between day 600 organoids and reference subclasses. Data points represent the average correlation value between day 600 organoids and clusters of a given subclass. **e** eTrace analysis of genes preferentially expressed in day 600 organoids. Squares represent query samples mapped to the eTrace and colored by the best-matching subclass (top annotation score). Point coordinates are based on similarity with reference clusters. Black arrows highlight apparent trajectories according to differentiation days. **f** 2D-UMAP representation of the single-cell RNAseq example of cerebral organoids^51^ colored by cluster (middle) and by best-matching transferred annotations (right). The barplot on the right represents transferred annotation subclasses by organoid time point. Data shows the proportion of cells in each organoid stage with a specific subclass annotation. **g** Expression values of known genes preferentially expressed in eIN (top-left), cRG (top-right), miEN (bottom-left), or imNIT (bottom-right) subclasses projected onto the 2D-UMAP representation of the organoid data. **h** Expression patterns of selected reference GO:BPs across organoid query clusters (right). Bottom annotations represent transferred subclass annotations (top), the organoid time point with the highest enrichment in the cluster (middle) and transferred stage annotations (bottom). The barplot on the right represents the maximum absolute expression of each GO:BP across organoid query clusters. Preferential expression scores of GO:BPs in corticogenesis network subclasses are used as reference (left).

The single-cell data includes iPSC- and embryonic stem cell (ESC)-derived cells profiled at different time points during cerebral organoid differentiation from pluripotency^51^. We pre-processed, clustered, and mapped the full dataset onto the corticogenesis network for interpretation (Methods) (**Fig. 6f**). We annotated each organoid cluster with best-matching reference subclasses based on network neighborhood. The relationship between the subclass annotations and the stage of differentiation recovers global biological expectation: early stages (IPSCs, EB) matched cycling radial glia cells, stages at 1 month of differentiation (1m) non-cycling radial glia, and subsequent stages (2-4m) immature excitatory and inhibitory neurons (**Fig. 6f**). The expression patterns of genes preferentially expressed in RGs or immature neurons *in vivo* overlapped with neighborhoods annotated with matching labels in a 2D single-cell landscape of the organoids, further supporting mapping and annotation transfer (**Fig. 6g**). To further interpret changes in molecular processes as organoids mature, we analysed the expression of selected signature biological processes of the developing cortex across organoid clusters (Methods) (**Fig. 6h**). The most active biological processes are consistent with subclass interpretations. For example, clusters annotated as cRG show high expression of mitosis-related processes (mitotic spindle checkpoint signaling, mitotic DNA replication), while clusters annotated as eIN express genes implicated in neurotransmitter transport and metabolism. Within the larger set of RG-matching cells, a subset of clusters show signatures of glial that match organoid maturation, suggesting presence of glial differentiation states in the more mature cells within the group. High expression of genes involved in stress response (e.g., heat, hypoxia) can be indicative of cellular stress^52,53^. Small clusters (e.g., c2, c20, c24) with high relative expression of genes associated with oxidative stress and unfolded protein response, together with joint elevated expression of processes commonly seen separately in neuronal or glial cells (e.g., synaptic signaling, lipid metabolism, and microglial activation) could be suggestive of potential cellular stress states.

Finally, we evaluated the quantitative metrics accompanying NeuRoDev’s mapping strategy using a wide range of comparisons with external and synthetic transcriptomics data, including scRNAseq data from 31 different tissues of the Human Protein Atlas (HPA) project^54^, from three human prefrontal cortex studies^12,17,55^ and randomly generated data (Methods). In each case, we mapped the data to the corticogenesis network and computed confidence scores measuring similarity between query samples and reference clusters. Data profiling the brain showed significantly higher confidence scores (mean=0.63) compared to other tissues (mean=0.24), or to randomly generated data (confidence=0.07) (**Supplementary Fig. S6c**; **Supplementary Table S4**), demonstrating the specificity of the metric and the mapping analysis. To evaluate the accuracy of the annotation transfer strategy, we mapped and annotated 11 pseudobulk samples generated *in silico* by aggregating cells from a scRNA-seq reference^21^ while controlling the ratio of neurons to radial glia (Methods) (**Supplementary Fig. S6d; Supplementary Table S4**). The annotation scores for neuronal subclasses highly correlated with the proportion of neuronal cells (cor=0.95; Pearson’s correlation). The scores for radial glia subclasses correlated with the fraction of radial glia cells (cor=0.95; Pearson’s correlation). To evaluate the ability of the mapping approach to detect relative differences in the composition of query samples, we generated 24 additional sets of pseudobulk profiles by combining single-cell transcriptomes extracted from an independent scRNAseq PFC dataset^55^. For each profile, we enriched one of the 24 cell types by doubling its cell number relative to the baseline composition. We then mapped their relative expression on the corticogenesis network and annotated each profile (Methods). Most annotations matched the expected (enriched) subclass, with few exceptions, suggesting that the mapping approach is able to detect differences in composition among query samples (**Supplementary Fig. S6e; Supplementary Table S4**). The annotation transfer strategy and accompanying metrics can thus accurately identify the most similar subclasses and detect relative differences between samples. Taken together, these analyses support the use of NeuRoDev to annotate and interpret query transcriptomic data. The mapping strategy recovers expected cell type and developmental stage annotations and provides resources to explore the molecular processes characterizing query samples.

## DISCUSSION

The availability of single-cell transcriptomic atlases profiling the human cortex at multiple developmental windows^11–21^ opens many opportunities to investigate the cellular biology of a long, protracted, and experimentally inaccessible process^5^. However, the use of such disparate data in daily practice is not straightforward. To contribute to this gap, from data generation to general accessibility and practical use, we developed NeuRoDev, a computational resource for the practicing neuroscientist interested in directly investigating the molecular and cellular processes unfolding during human cortical development. NeuRoDev combines curated data resources and novel analytical tools that facilitate the study of temporal and cellular patterns of gene expression in neuronal and glial cells as the human brain develops. It provides a reference framework to organize and interrogate this information, contextualize gene function, and interpret human experimental models.

Recent integration efforts aim to provide large brain cell references and expression signatures by assembling single-cell transcriptomes from cell atlases^52,56^, or identifying gene modules across many conditions^57^. We found that the integration and careful interpretation of clusters of cells identified independently in individual samples offers an intuitive alternative for integrative data analysis. NeuRoDev operationalizes this approach in the form of integrative summary networks. Networks offer a powerful framework to map and study the structure of complex biological systems^58,59^, and to represent and analyze heterogeneous data^60–63^. The integrative summary networks in NeuRoDev link clusters of cells that have similar transcriptional signatures and are reproducible across studies, thus facilitating robust data integration and analysis across developmental stages and data sources. The networks are accompanied by high-quality pseudobulk transcriptomic profiles that enable direct comparison of expression levels throughout the network and across individual clusters, cell subclasses, and developmental stages. This level of integration and scale allowed us to develop intuitive data analysis tools to directly explore cellular and temporal variability in developing neuronal and glia cells.

The eTrace analysis introduced herein maps expression patterns of individual genes or groups of genes in time, while tracking their identities and inferring developmental trends across a wide age range (from 5 weeks after conception up to 65+ years). As far as we know, there is no comparable analytical resource available to easily interrogate these patterns in human data. We anticipate that this tool will be useful in supporting hypothesis generation when studying gene function, cellular maturation, and patterns of convergence of disease-associated variants, or when selecting the relevant physiological context in which to study the effects of loss-of-function mutations in neurodevelopmental disease genes^63–65^. The original data and reference networks in NeuRoDev are highly curated. Although we define cell subclasses to aid interpretation, the study of cell variability in NeuRoDev need not be restricted to these predefined groups. We expect additional meaningful cellular states to be represented in our networks without being labeled, and such patterns can be interrogated directly by analysing and comparing pseudobulk profiles. eTrace analysis also facilitates this unbiased analysis by allowing direct observation of all individual clusters.

Because of the known differences in human neurodevelopment relative to other species, a full understanding of how the human brain develops and functions requires direct analysis of human cells^5^. *In vitro* human experimental systems are an important development in this regard. However, there are limitations to the level of maturation achieved by these systems. A more thorough understanding of the processes underlying human brain development *in vivo* is needed to answer fundamental questions in biology and medicine and better recapitulate human neurodevelopmental processes in model systems. Our resource contributes to this need by providing a means to inspect molecular and cellular processes as they unfold in human tissue. NeuRoDev is provided as a data resource, a software package, and web applications for interactive data visualizations and exploration.

## METHODS

### Data sources

**Single-cell transcriptomics data compendium** snRNAseq data profiling human fetal or adult cortical samples were compiled from 11 independent studies^11–18,20,21^, for a total of 1,617,236 single-cell transcriptomes. The selected studies collectively survey all stages of development, from post-conception week 5 to 64 years of age (**Supplementary Table 1**).

**Reference genes** Gene Ontology biological processes (GO:BP), molecular functions (GO:MF), and cellular components (GO:CC) were obtained from the *EnrichR*^66^ database (available at https://maayanlab.cloud/Enrichr/#libraries, release 2025). Only gene sets that contained at least five genes present in our resource were considered for further analysis (GO:BP nSets=4,344, nGenes=11,265; GO:MF nSets=924, nGenes=8,636; GO:CC nSets=396, nGenes=9,141). Neuronal GO:BP (nSets=79, nGenes=2,702) were selected based on the word “neuronal” in the gene sets annotations, keeping genes with an average expression across neurogenesis of at least log2(CPM)=1. Ion channel families (239 genes, 29 families) were downloaded from *The Guide to PHARMACOLOGY database* (https://www.guidetopharmacology.org/download.jsp) (targets_and_families_v2025.2). HCN channels originally included within Cyclic nucleotide-regulated channels (CNG) channels were classified as a separate family.

#### Single-cell compendium preprocessing

**Proportional sampling** single-cell level data analyses and clustering was performed independently per sample. To obtain balanced samples, each dataset was first divided based on age to obtain 119 samples. To better balance cell number, 23 samples with a number of cells higher than the average across all samples (∼14,000) were proportionally downsampled. To this end, each outlier sample was first clustered using the standard clustering pipeline of the R package Seurat (https://satijalab.org/seurat/) and then downsampled to ∼14,000 cells by randomly subsampling cells from each cluster maintaining the original cluster proportions of the full sample (see subsample clustering). This process resulted in a total of 1,001,928 cells. For simplicity, following this preprocessing step we refer to all samples as subsamples.

**Consensus labeling** Individual cells in all subsamples were annotated with consensus class and subclass labels selected from 3 cell classes (*glial, neuronal, other)* and 11 cell subclasses (*RG = radial glia, IP = intermediate progenitor, OPC = oligodendrocyte progenitor cell, Ast = astrocyte, Oli = oligodendrocyte, Mic = microglia, ExN = excitatory neuron, InN = inhibitory neuron, Imm = immune cell, Vas = vascular, Unk = unknown*). These consensus annotations were manually generated based on those reported in the original studies. Cells were also annotated with consensus age and developmental stage labels. Age labels were defined using post-conceptional week (*pcw*) units for prenatal subsamples, and days (*d*), months (*m*), or years (*yr*) for postnatal subsamples. A total of 79 unique age labels were defined. Developmental stage labels (n=13) were defined based on Brain Span project guidelines (https://www.brainspan.org/).

### Network-based summary data integration

**Subsample clustering** Single-cell clustering was performed individually per subsample in all cases using the standard Seurat pipeline (https://satijalab.org/seurat/) (*NormalizeData*, *FindVariableFeatures*, *ScaleData*, *RunPCA*, *FindNeighbors*, *FindClusters*) with number of dimensions (*dims=1:10)* in *FindNeighbors,* resolution=*0.5* in *FindClusters* and otherwise default parameter values.

**Network-based integration** To integrate data across subsamples and studies, clusters were first represented based on their transcriptional similarity to a panel of cell type references. To this end, a transcriptional profile was defined per cluster by averaging the normalized and log2-scaled count data across the cells belonging to the cluster. These profiles were used to define a relative expression profile per cluster (cluster signature) by centering the profile relative to the average profile across clusters in the same subsample. This procedure resulted in cluster expression signatures that remove common subsample-dependent expression patterns. These signatures were then correlated with equivalent transcriptional signatures obtained by comparing reference cell type profiles (see below) resulting in transcriptional cell-type similarity representation (i.e., a vector of Pearson’s correlation coefficients per cluster estimating similarity to predefined cell type signatures). The cell-type similarity representation was used to measure similarity between clusters and build a network connecting them accordingly. The Euclidean distance was used as a similarity metric. To define a final pruned and weighted network connecting clusters, the Shared Nearest Neighbor (SNN) algorithm^67^ was used considering the 15 closest neighbors of each cluster. The network is weighted considering both the absolute number of shared neighbors between cluster pairs and their rank similarity. For network layout and visualization purposes, each node was assigned coordinates based on the UMAP^68^ algorithm as implemented in the *umap* R package (https://CRAN.R-project.org/package=umap) using cell-type similarity representation as input (*n_neighbors=15*).

**Reference cell type signatures** Cell type reference signatures were defined based on the full data and reported cell type annotations of 6 reference studies^11,12,14,17,20,21^. These studies were selected based on developmental stage coverage and cell label interpretability. Cell type signatures were estimated for all annotated cell types within each independent dataset following the same procedure used for network-based integration. For lineage-specific networks, only lineage-specific cell types were considered, and signatures were computed per dataset relative only to the subset of lineage-specific cell types. A total of 159, 106, and 62 cell type signatures were obtained respectively for the corticogenesis, neurogenesis, and gliogenesis networks.

**Cluster annotation** Clusters were interpreted in terms of putative cell identities and states based on multiple sources of evidence, including transcriptional similarity with reference cell types, marker gene expression, network neighborhood, age, and developmental relationships. For simplicity, we refer to the resulting labels as subclasses. A preliminary set of candidate cell type labels was first estimated based on best-matching cell types according to the correlation values between transcriptional signatures of clusters and reference cell types. To reduce label redundancy and merge similar cell types, reference networks were then clustered using the *Leiden* clustering algorithm as implemented in the R package *igraph (*https://cran.r-project.org/package=igraph). A resolution parameter value of 1 was used for the corticogenesis network, and a value of 0.75 for the lineage-specific networks. Subsequently, the network topology was leveraged to further infer coherent labels following a majority-voting algorithm. Each original cluster was iteratively re-annotated based on the labels of its neighbors. Labels were updated based on the highest frequency of neighbor labels. In cases of equal frequency, the label with the highest edge weight average was assigned. The procedure was repeated until reaching stability, as determined by 10 iterations of unchanging labels for 99% of clusters. Because of differences in network size and subclass resolution, different numbers of neighbors were considered for corticogenesis (n=25) and lineage-specific networks (n=5). These numbers were determined empirically by testing different values. Patterns of marker gene expression based on pseudobulk profiles were subsequently used to assess consistency with estimated labels and expected cell type expression. Cluster labels with inconsistent evidence were inspected further by considering the correlation values with all reference cell types, their top signature genes, and the consistency between putative cell types and developmental stages. Inconsistent clusters underwent further quality checks (see *Quality control*). The final 21 (corticogenesis), 25 (neurogenesis), 18 (gliogenesis) putative subclasses reported (**Supplementary Table S3**) passed all quality checks and manual inspection.

**Quality control (QC)** For clusters with inconsistent annotation labels, the following parameters were further inspected: the number of detected genes, the total number of counts, the percentage of ribosomal genes, and the percentage of mitochondrial genes. Clusters were considered of poor quality if they showed values significantly lower or significantly higher compared to the distribution of all clusters. 1575, 1248, and 785 clusters passed all QC steps in the corticogenesis, neurogenesis, and gliogenesis networks. Low quality clusters with similar properties were robustly identified across datasets (data not shown), highlighting an additional useful feature of the integrative approach proposed herein. Based on these QC procedures, corresponding single-cell transcriptomes were also filtered-out from the final compendium. The total number of single-cell transcriptomes that passed preprocessing and quality control and whose profiles are summarized in at least one reference network is 948,350. The final corticogenesis network summarizes a total of 930,455 single-cell transcriptomes; the neurogenesis network 556,045; and the gliogenesis network 240,737.

**Pseudobulk transcriptomes** Each reference network in NeuRoDev is accompanied by a set of high-quality transcriptional pseudobulk profiles (one per cluster; 1575 corticogenesis, 1248 neurogenesis, and 785 gliogenesis). A pseudobulk transcriptome was generated for each cluster by aggregating the total read count across all cells in the cluster and removing study-associated batch effects, while considering stage and subclass annotations as biological covariates. The *ComBat_seq* function of the R package *sva* (https://bioconductor.org/packages/sva) was used for count-level batch correction. Corrected counts were normalized as Counts Per Million (CPM) and log2-scaled.

### Differential expression analyses

**Preferential expression** Preferential gene expression was quantified by performing pairwise differential expression analyses between subclass pseudobulk profiles using linear models as implemented in the R package *limma (*https://bioconductor.org/packages/limma/). Expression values were first normalized using the *calcNormFactors* function of the R package *edgeR (*https://bioconductor.org/packages/edgeR/). A single preferential expression score per gene and subclass was defined as the sum of the differential expression coefficients in all pairwise comparisons between that subclass and all other subclasses. A large and positive score indicates higher expression in that subclass relative to each of the other subclasses, with the magnitude of the value quantifying how large the difference is. Only significant coefficients at a Bonferroni-adjusted p-value < 0.01 were considered for the aggregate score. The final aggregate score was normalized by the total number of subclasses.

**Differential expression between conditions** All differential expression analyses between two groups were performed by fitting linear models as implemented in the R package *limma*. Differential expression mature and developing excitatory neurons (**Fig. 2**) compared the groups: mature (clusters belonging to the L6IT, L6NIT, L35IT, and L23IT subclasses; n=505) versus developing (clusters belonging to the miEN, imNIT, eEN, and imIT subclasses; n=167).

**Rank-based gene set enrichment analysis** All enrichment analyses were performed using the R package *fgsea* (https://bioconductor.org/packages/fgsea/) to estimate overrepresentation or query genes within extreme ranking values.

**Preferential gene groups** Genes were assigned to a subclass or set of subclasses (subclass group) based on the preferential expression scores for individual subclasses. Genes were first assigned to one or more subclasses if their preferential score for the subclass was larger than a predefined threshold (>1.5). This threshold was determined empirically based on the distribution of all preferential expression values. Selected genes were further required to meet two additional criteria: a log2 fold-change > 2 when comparing the average expression in the subclass/group where they were assigned to versus the average expression across the rest of subclasses, and an average preferential expression score > 2 across all individual scores corresponding to the subclasses forming the group.

**Inferred temporal expression trends** Inferred expression trends were computed by fitting a spline function between time steps and query values as implemented in the *smooth.spline* function of the R package *stats*. For the case of gene sets (e.g., GOBP processes) a “pathway” expression score was first estimated via Gene Set Variation Analysis based on pseudobulk transcriptomes as implemented in the R package *GSVA* (https://bioconductor.org/packages/GSVA/) (*gsva* function) and then fitted with splines.

**Preferential expression of GO processes** Preferential expression values and sets for GO:BP/MF/CC genesets were defined in the same way as for individual genes but using “pathway” expression score profiles instead of gene expression values. The resulting scores were processed following the same procedure described in *Preferential expression* but adjusting parameters to consider prenormalized values as input. GO processes were assigned to subclass groups following the same steps described in *Preferential gene groups,* except for the final filtering steps. Only groups with at least 2 processes were considered.

**Selection of representative GO signature processes** A set of Gene Ontology Biological Processes (GO:BP) was selected to aid the interpretation of cellular states in reference networks and query data. Processes were selected manually according to preferential expression patterns to represent 10 major interpretable groups: RG, immEN, matEN, matIN, Astro, Micro, OligoGlia, Stress (**Supplementary Fig. S6h**). Each group includes a set of GO:BPs (n∼4) that are preferentially expressed in the corresponding subclasses and physiologically interpretable. The stress group was defined based on function and is intended to aid interpretation of external data and flag potential artefacts. An additional set of GO:BPs (n=20) was selected to represent the neuronal lineage based on the neurogenesis network and known differentiation/maturation processes.

### eTrace analysis

**eTrace** To visualize patterns of expression variability across cell subclasses and time with a simple plot, we introduce eTrace analysis. An eTrace plots a given molecular phenotype score estimated from the pseudobulk transcriptomes of one of the reference networks (y-axis) as a function of time/age (x-axis). Each value shown in the plot corresponds to one of the clusters in the network. Clusters are colored based on their corresponding developmental stage and cellular subclass. The default molecular phenotype shown is gene expression enrichment (see *Expression enrichment*).

**Expression enrichment** For eTrace analysis, gene expression enrichment can be estimated for individual genes or gene groups. For individual genes, enrichment is measured by the deviation of the gene’s expression level from the average expression across all clusters scaled by its standard deviation (z-scale). This value is plotted by default. For gene groups, enrichment is measured by the z-score statistic estimated by comparing the observed average relative expression (z-scale) of query genes with that expected from a null distribution of randomly sampled gene sets of equal size. The z-score is plotted by default. Relative expression values, z-score statistics, and corresponding p-values are computed and reported. These calculations are implemented in the *get_eTrace* and *plot_eTrace* functions in the neuRoDev R package.

**Stage-specific subclass enrichment** Expression enrichment scores for all pairs of developmental stages and subclasses are estimated as follows pairwise by subsetting the corresponding pseudobulk expression profiles, z-scaling expression values across all clusters in the subset, estimating the average scaled expression value of query genes, comparing these values with random expectation from randomly sampled gene sets of equal size to estimate a z-score statistic and corresponding p-value. Enrichment scores measured by these z-scores are plotted in matrix format with subclasses in the x-axis and developmental stages in the y-axis in developmental order from top to bottom. These calculations are implemented in the *get_eMatrix* and *plot_eMatrix* functions in the neuRoDev R package.

### Query data mapping and interpretation

**Network mapping and visualization** The mapping of query transcriptomic data onto a reference network is based on the similarity between input profiles and the reference pseudobulk profiles of the clusters in the network as measured by the Pearson’s correlation coefficient. A preselected subset of informative genes is used as features for computing the correlation. Informative genes (n∼500) were selected based on top preferential expression values per subclass. Query samples are projected on the same 2D coordinate system of the reference network by estimating for each sample a pulling score based on the transcriptional similarity and 2D coordinates of its closest neighboring nodes in the network (n=15 by default). Closeness (similarity) is defined based on correlation values. The pulling score represents an attractive force towards a reference cluster in the 2D coordinate system. The pulling score is computed as a network-centered weighted sum of the coordinates of closest neighbors. The coordinates are weighted by correlation values scaled between 0 and 1. The weighted sum is centered by adding the center of the reference network coordinates. The resulting 2D coordinate represents the pulling scores for the x and y coordinates. High correlation values with similarly located clusters will pull a query point towards those clusters. Low correlation values with sparsely located clusters will place the point in the center of the clusters. For query scRNAseq data, input single cell data is first clustered to generate pseudobulk profiles for query clusters. These profiles are then mapped following the correlation approach. Query bulk RNAseq data are correlated directly.

**Mapping confidence score** To quantify how well a query dataset maps to a reference network, we introduce two types of confidence scores. A sample score is assigned to each query sample. A global score is defined for the totality of query data. Sample confidence score is measured as the average correlation value between the query transcriptional profile and the pseudobulk transcriptional profiles of the nearest neighbor clusters (n=15 by default) in the target network. Correlation is measured by the Pearson’s correlation coefficient computed using only the preselected set of informative genes as features. Global confidence score is measured by the average sample score estimated across all query samples mapped.

**Reference knowledge scoring for query data interpretation** To aid the biological interpretation of query samples mapped to a reference network, we introduced similarity measures indicating what fraction of the transcriptionally similar neighborhood of a query sample corresponds to a given label (e.g., cell subclass or developmental stage). For each query sample mapped to the network, a single score per individual reference label is measured based on the annotations of the nearest neighboring clusters (n=15 by default). An annotation similarity score per unique label is measured as the sum of the correlation values corresponding to neighboring clusters labeled with that label. Label scores are normalized by the total sum across labels to derive fractional scores. Thus, a single query sample is scored with a vector of numbers, one number for each unique label present in the neighborhood.

**eTrace mapping** Query data is mapped onto eTraces by calculating 2D coordinates using the average coordinates of top 15 most correlated reference pseudobulk profiles.

**Query data interpretation with GO signature processes** Expression of manually curated Gene Ontology Biological Processes (GO:BP) in query datasets was visualized by taking the average expression of genes in each GO:BP for each sample/cluster (mapped point) and then scaled by subtracting the mean and dividing by the standard deviation across mapped points. The maximum absolute average expression (not scaled) of each GO:BP across mapped points was computed and shown as a barplot.

### Organoid query data analyses

**Organoid bulk RNAseq mapping analysis** Published data from cortical organoids profiled with bulk RNAseq at 11 different time points (days 25, 50, 75, 100, 150, 200, 250, 300, 400, 500, 600; total replicates=62)^50^ were preprocessed and mapped onto the corticogenesis reference network. Prior to mapping, a batch correction step was performed using the R package *sva* and reported batch labels, and the data normalized as log_2_ counts per million after correction. Average samples across replicates per time point were used as query profiles for mapping. Differential expression analysis between time points was performed by comparing expression levels of at each time point versus all others using the R package *edgeR*.

**Organoid single-cell RNAseq mapping analysis** Published data from cerebral organoids profiled with single-cell RNAseq at 7 differentiation stages (iPSC, embryoid body, neuroectoderm, neuroepithelium, 1 month, 2 months, 4 months; total cells=90,576)^51^ were clustered and mapped on the corticogenesis network. Cells annotated as “Glyc” were not considered for the analysis (remaining cells=75,265). Expression normalization and clustering was performed using the R package *ACTIONet*^61^ with the following functions: *normalize.ace*, *reduce.ace*, *runACTIONet*, *clusterCells*. For clustering the resolution parameter was set to 2. Default values were used for all other parameters.

### *In silico* evaluation

**Confidence score evaluation** To test the applicability of the confidence score in multiple settings, scRNAseq data from 31 different tissues from the Human Protein Atlas^54^ project were mapped onto the corticogenesis reference network (see: **Supplementary Table S4**). The query tissues include adipose tissue, bone marrow, brain, breast, bronchus, colon, endometrium, esophagus, eye, fallopian tube, heart muscle, kidney, liver, lung, lymph node, ovary, pancreas, pmbc (peripheral mononuclear blood cells), placenta, prostate, rectum, salivary gland, skeletal muscle, skin, small intestine, spleen, stomach, testis, thymus, tongue, and vascular. For each tissue, the originally reported clusters were used for mapping. Normalized expression profiles (nTPM) of clusters annotated with the same cell type were averaged to define a single expression profile for each annotated cell type. These cell types were used as input query samples and mapped independently per tissue. One additional independent scRNAseq dataset profiling the human prefrontal cortex of healthy individuals was obtained from^55^ and processed to obtain average normalized transcriptional profiles for each originally annotated cell type. Two additional reference datasets also included in our resource ^17, 12^ were processed to obtain transcriptional profiles for reported cell types. Finally, for contrast, a random dataset was generated using a uniform distribution with values between 0 and 5 and included in the comparison. All query samples from the different datasets were mapped and scored using the proposed confidence score metric (see mapping section above). Confidence scores were ranked for comparison, as shown in **Supplementary Fig. S6c**.

**Reference knowledge scoring evaluation** To test the mapping annotation when using transcriptional profiles, eleven pseudobulk profiles were computed from a reference scRNAseq dataset^21^ as follows. In each case, 2000 cells per pseudobulk were randomly selected, each time increasing the percentage of neuronal cells considered (annotated as EN-L6-CT) while reducing the percentage of radial glia cells (annotated as RG-vRG). Proportion changes were considered from 0% to 100% with a 10% step. The resulting pseudobulks were mapped on the corticogenesis network and scored based on neighboring reference labels. The procedure was repeated 20 times to reduce biases arising from random sampling, and the annotation scores were averaged across iterations. Based on these scores, overall neuronal and glial scores were estimated. A neuronal score was estimated as the sum of the annotation scores for the mature neuron subclasses (L23IT, L35IT, L6IT, L6NIT), a radial glia score was as the sum of the scores for RG and cRG subclasses. These overall scores were correlated (Pearson’s correlation) with the original known radial glia to neuron ratios to evaluate how well the annotation captures the changes in proportion. The resulting annotation scores are reported in **Supplementary Table S4**.

**Relative composition evaluation** To evaluate whether the mapping approach detects changes in relative differences between samples, 24 pseudobulk profile sets were generated by combining in controlled proportions single-cell transcriptomes extracted from an independent scRNAseq dataset^55^. In each case the proportion of one cell type was doubled while adjusting the proportions of others to keep the total sum to 1. Next, 25,000 cells were randomly sampled while keeping the predefined proportion structure and their expression counts were summed and normalized to obtain a pseudobulk profile per cell type. The relative expression signatures (row centered) of the obtained pseudobulks were then used as query input for mapping. This procedure was repeated 20 times per test proportion case to account for biases arising from random sampling. The annotation scores were averaged across the 20 iterations. The annotation score corresponding to the cell type whose proportion was doubled was compared with the known proportion. The annotation scores are reported in **Supplementary Table S4**.

## DATA AVAILABILITY

The data that support the findings of this study are available in Figshare and the Supplementary Data files. All previously published reference transcriptomic data are publicly available with sources listed in Supplementary Table S1. Full processed data of the curated compendium and integrative networks are available at Figshare with accession https://doi.org/10.6084/m9.figshare.30885428.

## CODE AVAILABILITY

The neuRoDev R package and a tutorial to reproduce analyses are available at https://github.com/davilavelderrainlab/neuRoDev. Interactive web applications for eTrace analysis in cortical, neuronal, and glial development are available at: https://erikbot.shinyapps.io/etraceshinycortico/, https://erikbot.shinyapps.io/etraceshinyneuro/, and https://erikbot.shinyapps.io/etraceshinyglio/.

## ACKNOWLEDGEMENTS

We thank members of the Davila-Velderrain group for discussion and feedback. This research was supported by the Human Technopole Foundation. A.Z and E.B are PhD students within the European School of Molecular Medicine (SEMM).

## AUTHOR CONTRIBUTIONS

A.Z, E.B, and J.D.V performed computational analyses. E.B implemented the R package with help from A.Z. All authors conceptualized the study, designed and created the figures, and wrote the manuscript. J.D.V supervised the work.

## COMPETING INTERESTS

The authors declare no competing interests.

## Supplementary Figures

**Supplementary Figure S1.**
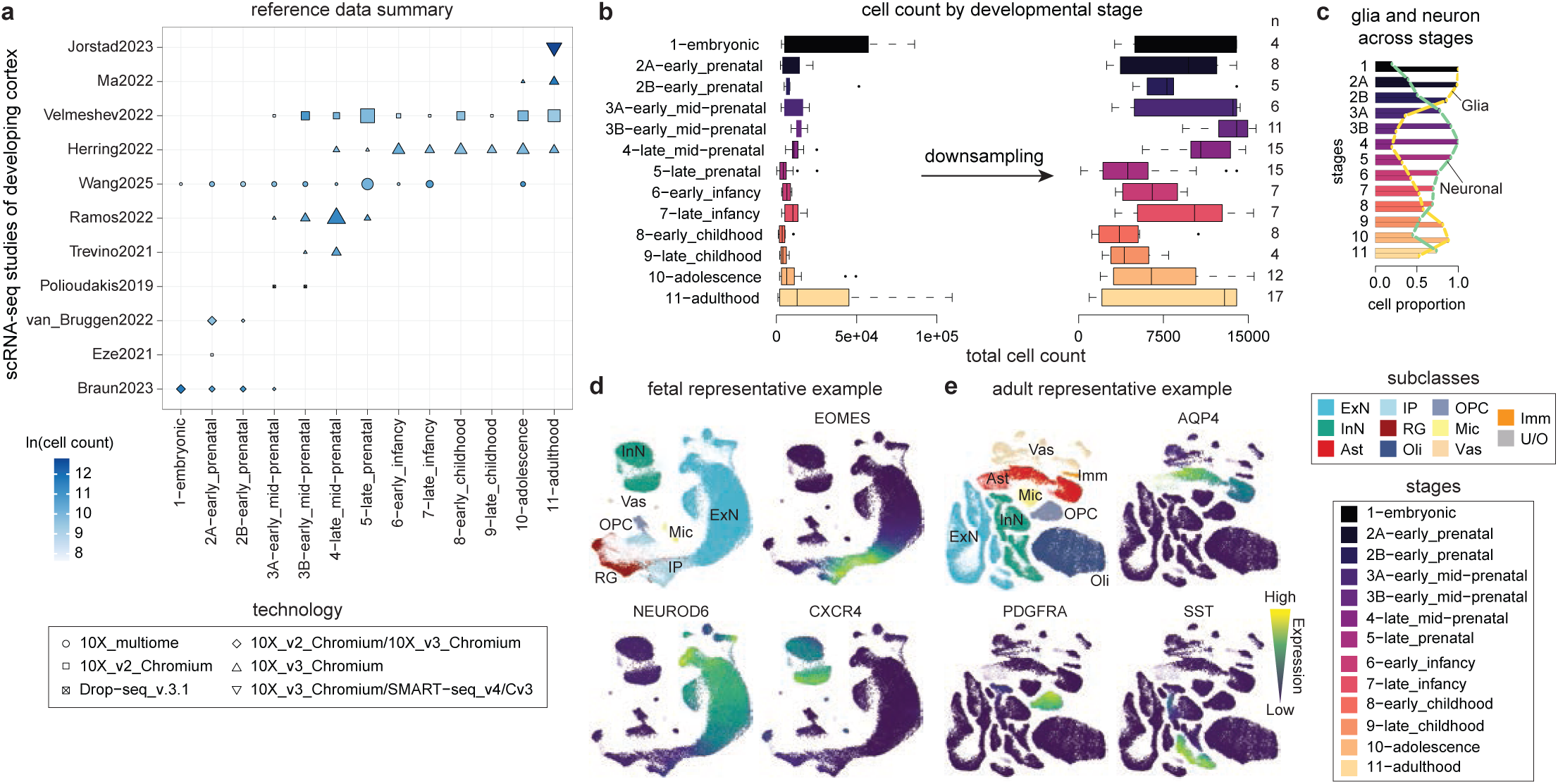
Single-cell cortical development compendium statistical summary and harmonization. **a** Summary information of published datasets compiled (rows) (reported as first author name-year of publication; n=11) over developmental stages (columns; n=13). Shapes indicate the scRNA-seq technology. Colors indicate the number of cells in log scale (11.40 ± 0.23 cells per stage; 11.36 ± 0.44 cells per dataset; 10.47 ± 0.23 cells per stage-dataset pair; mean ± s.e.m.). Size represents the number of samples (9.15 ± 1.22 samples per stage; 10.82 ± 3.26 samples per dataset; 2.59 ± 0.23 samples per stage-dataset pair; mean ± s.e.m.). **b** Total cell counts per developmental stage before (left; 1.24⋅10^5^ ± 3.52⋅10^4^ cells per stage; mean ± s.e.m.) and after (right; 7.71⋅10^4^ ± 1.35⋅10^4^ cells per stage; mean ± s.e.m.) downsampling. **c** Glia and neuronal cell composition across stages. The proportion is normalized for each class to its maximum across all stages (glia maximum = stage 1; neuronal maximum = stage 4). **d-e** 2D-UMAP representation of a single-cell landscape of fetal^12^ (d) and adult^17^ (e) representative datasets. Data points represent single cells colored by subclass (top-left) or by the level of expression of marker genes: EOMES (intermediate progenitors; top-right), NEUROD6 (developing excitatory neurons; bottom-left), CXCR4 (immature inhibitory neurons; bottom-right) (d), AQP4 (astrocytes; top-right), PDGFRA (OPC; bottom-left), and SST (mature inhibitory neurons; bottom-right) (e). Box-plots are centered around the median; the interquartile range (IQR) defines the box; whiskers extend to the largest(smallest) value no further than 1.5 × IQR from the end of the box.

**Supplementary Figure S2.**
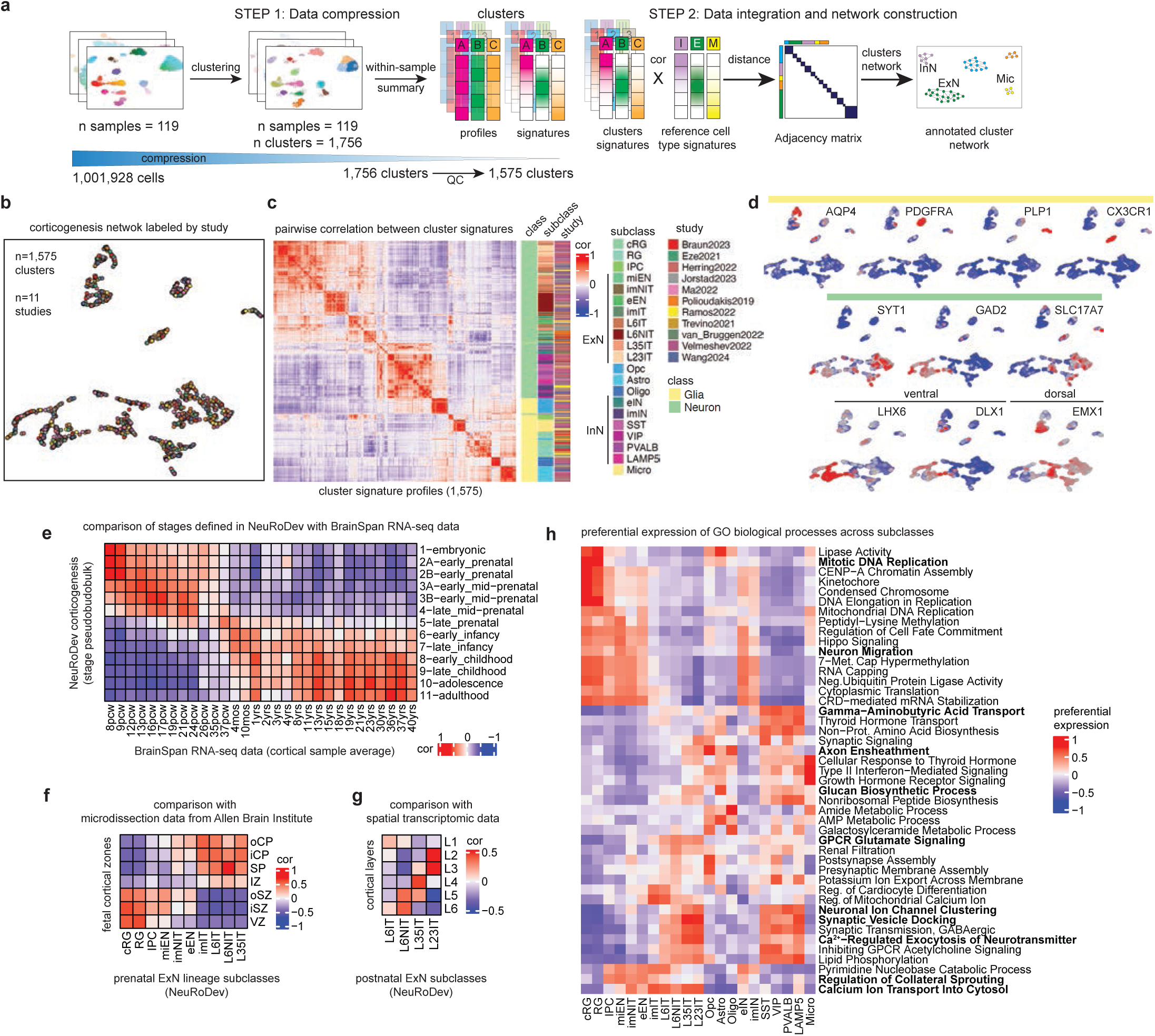
Overview of data integration, validation, and annotation analyses. **a** Schematic illustrating dataset compression and integration steps. **b** 2D-UMAP representation of the corticogenesis reference network colored by the study of origin. Data points represent single clusters (n=1,575). **c** Pairwise correlation between cluster signatures. Annotations on the right highlight the class, subclass, and study of each cluster. **d** Expression enrichment values of glial (top), neuronal (middle), ventral (bottom-left), and dorsal (bottom-right) marker genes across the corticogenesis reference network. Red = high, blue = low. **e** Comparison between developmental stages defined in NeuRoDev and BrainSpan RNA-seq data. Data shows the correlation between NeuRoDev stage pseudobulk profiles and cortical BrainSpan RNA-seq profiles averaged by age. **f** Comparison between NeuRoDev prenatal excitatory neuron lineage subclasses and microdissection data from the Allen Brain Institute^6^. Data shows the correlation between pseudobulk profiles of NeuRoDev subclasses and microdissection bulk RNA-seq profiles. **g** Comparison between NeuRoDev postnatal excitatory neuron lineage subclasses and spatial transcriptomics data^24^. Data shows the correlation between pseudobulk profiles of NeuRoDev subclasses and transcriptomes of corresponding cortical layers defined by spatial transcriptomics. **h** Preferential expression of Gene Ontology biological processes across subclasses. Top 3 preferentially expressed processes per subclass are shown.

**Supplementary Figure S3.**
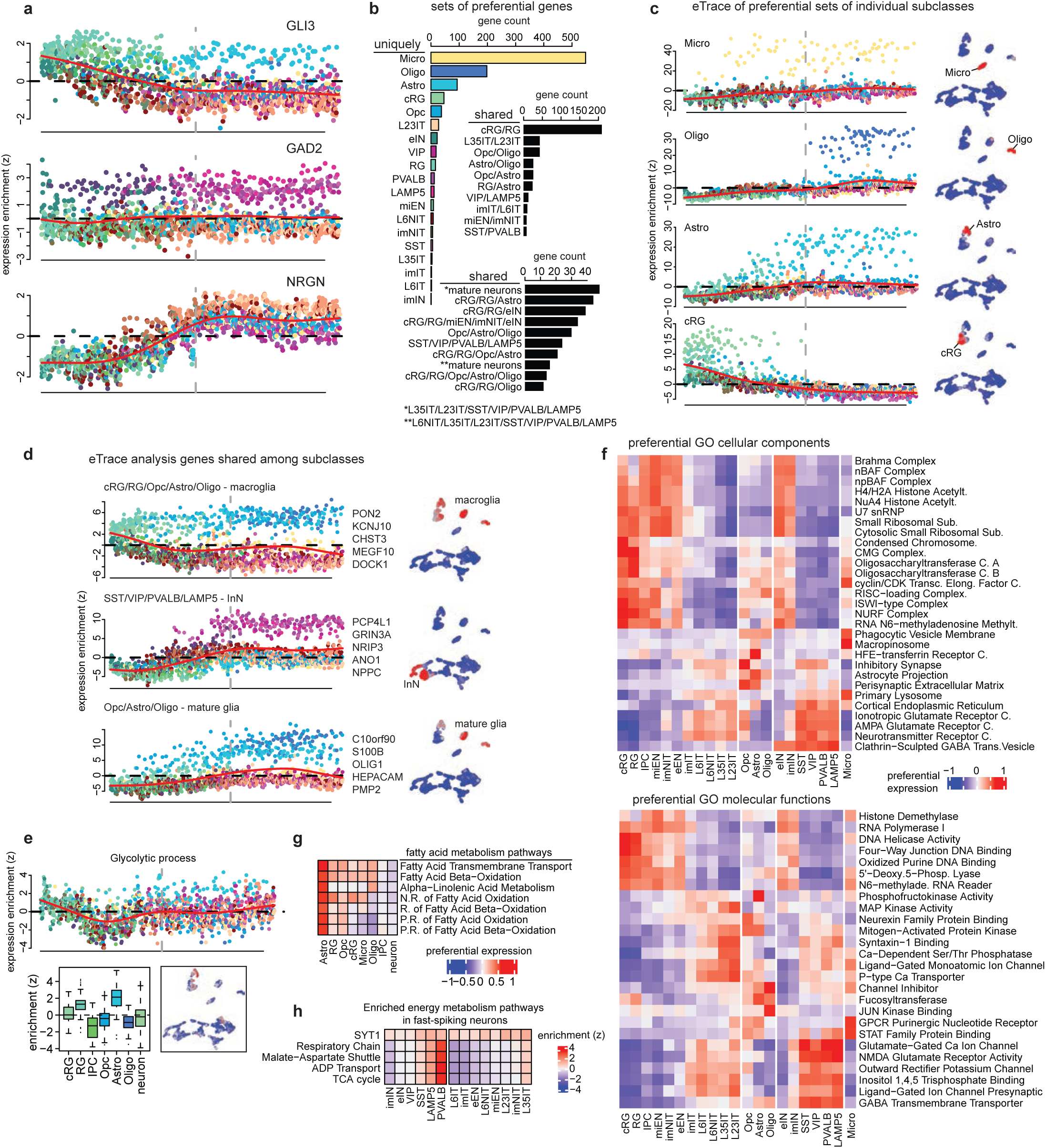
eTrace analysis of subclass exclusive and shared genes and biological processes. **a** eTrace analysis of marker genes GLI3 (RG, cRG, Astro), GAD2 (inhibitory neurons), and NRGN (mature excitatory neurons). Points represent single clusters (n=1,575). **b** Number of genes preferentially expressed in each subclass (colored barplot), in two subclasses (black barplot, top), and in more than two subclasses (black barplot, bottom). **c** eTrace analysis (left) and expression enrichment network projection (right) of genes preferentially expressed in microglia (Micro), oligodendrocytes (Oligo), astrocytes (Astro), and cycling radial glia (cRG) (right; red = high enrichment, blue = low enrichment). **d** Same as (**c**) for genes preferentially expressed in macroglia, mature inhibitory neurons, and mature macroglia. Five representative genes for each gene set are shown (middle). **e** Expression enrichment of genes involved in the glycolytic process shown as eTrace (top), distribution in progenitors (cRG, RG, IPC), opc (Opc), astrocytes (Astro), oligodendrocytes (Oligo), and neurons (bottom left), and across the corticogenesis network (bottom right). **f** Preferential expression of Gene Ontology cellular components (top) and molecular functions (bottom) across subclasses. The top-2 preferentially expressed sets are shown for each subclass. **g** Preferential expression of fatty acid metabolism pathways across glia (cRG, RG, IPC, Opc, Oligo, Astro, Micro) and neurons. **h** Expression enrichment of energy metabolism pathways that are significantly enriched in parvalbumin neurons (PVALB) across inhibitory and excitatory neurons. C. = complex.

**Supplementary Figure S4.**
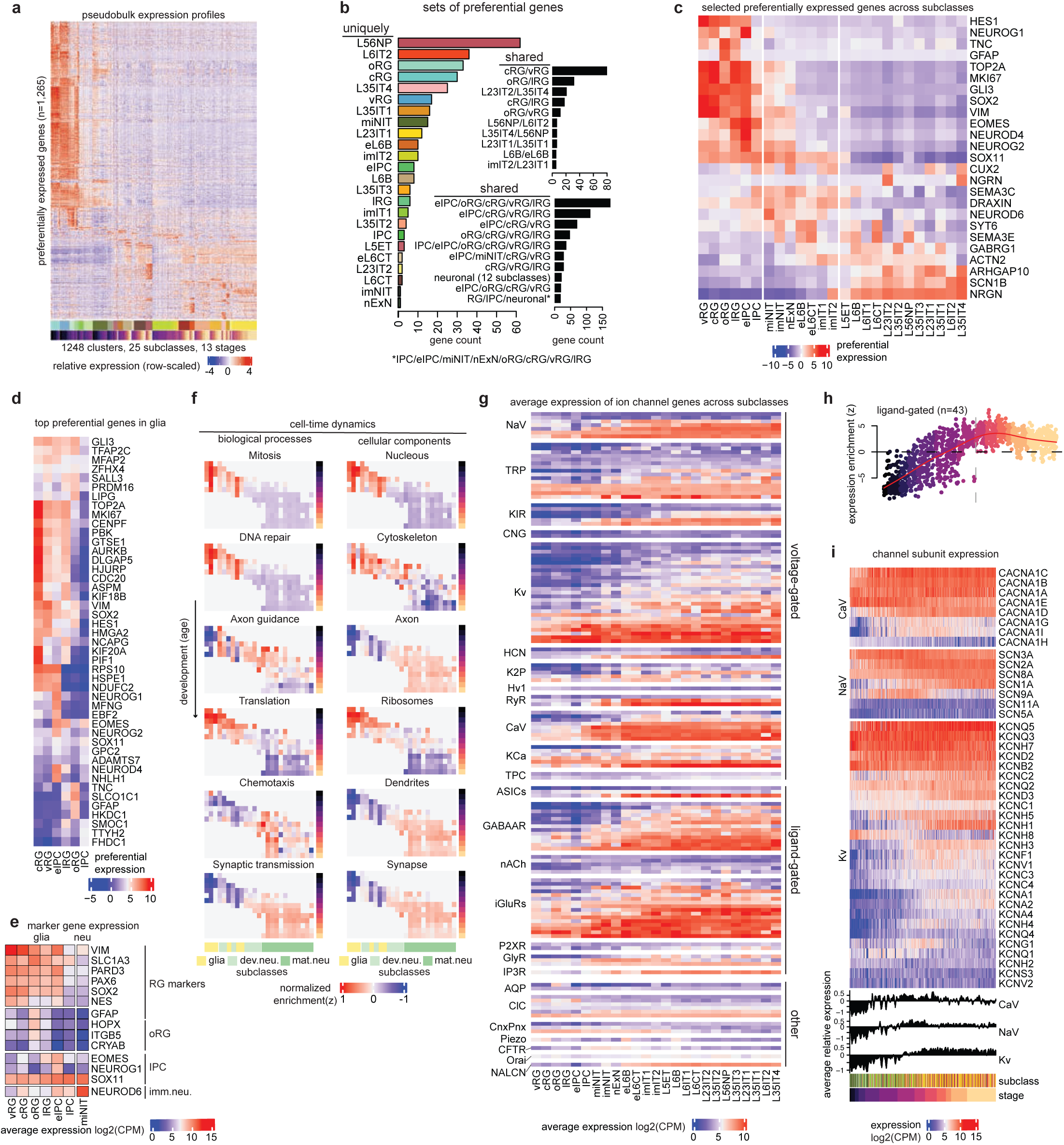
Structural and functional maturation patterns in neurogenesis network. **a** Relative expression (row z-scaled) of preferentially expressed genes across neurogenesis clusters. **b** Number of genes preferentially expressed in each subclass (colored barplot), in two subclasses (black barplot, top), and in more than two subclasses (black barplot, bottom). **c** Preferential expression of selected genes across subclasses. **d** Preferential expression of the top preferentially expressed genes in glial progenitors. **e** Average expression (log-normalized counts per million) of selected markers in radial glia (RG), outer RG (oRG), intermediate progenitor cells (IPC), and immature neurons. **f** Expression enrichment of selected Gene Ontology biological processes (left) and cellular components (right) across subclasses and developmental stages. **g** Expression (log-normalized counts per million) of ion channel genes across subclasses. Ion channel genes are classified by family. Subclasses are ordered based on average age. **h** eTrace analysis of ligand-gated ion channels colored by developmental stage. **i** Expression (log-normalized counts per million) of ion channel genes (voltage-gated calcium, sodium, and potassium channels families) across age-ordered clusters. Bottom barplots show average relative expression values across all genes of each family.

**Supplementary Figure S5.**
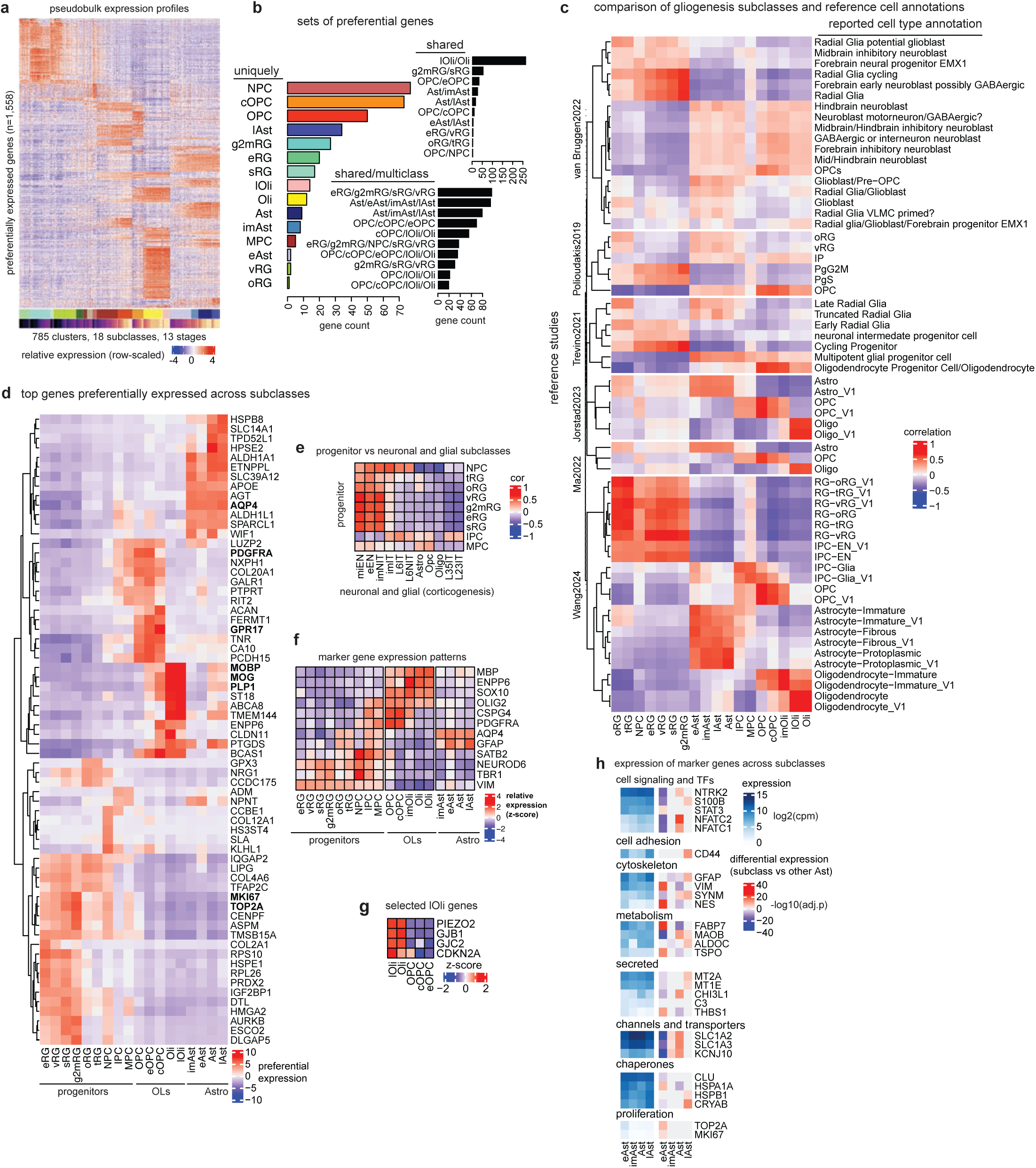
Expression signatures in glial development network. **a** Relative expression (row-scaled) of preferentially expressed genes across gliogenesis clusters. **b** Number of genes preferentially expressed in each subclass (colored barplot), in two subclasses (black barplot, top), and in more than two subclasses (black barplot, bottom). **c** Similarity between gliogenesis subclasses and reference cell annotations (Pearson’s correlation). **d** Preferential expression across gliogenesis subclasses. The top-5 most preferentially expressed genes per subclass are shown. **e** Similarity between progenitor subclasses in the gliogenesis network and neuronal and glial subclasses in the corticogenesis network (Pearson’s correlation). **f** Relative expression (row z-score) of selected marker genes across subclasses. **g** Relative expression (row z-score) of genes overexpressed in lOli in the oligodendroglia lineage. **h** Expression (log-normalized counts per million; left) and differential expression (-log_10_ of the adjusted p-value, one-vs-all differential expression analysis; right) of selected marker genes across the astroglia lineage. Genes are divided based on their function.

**Supplementary Figure S6.**
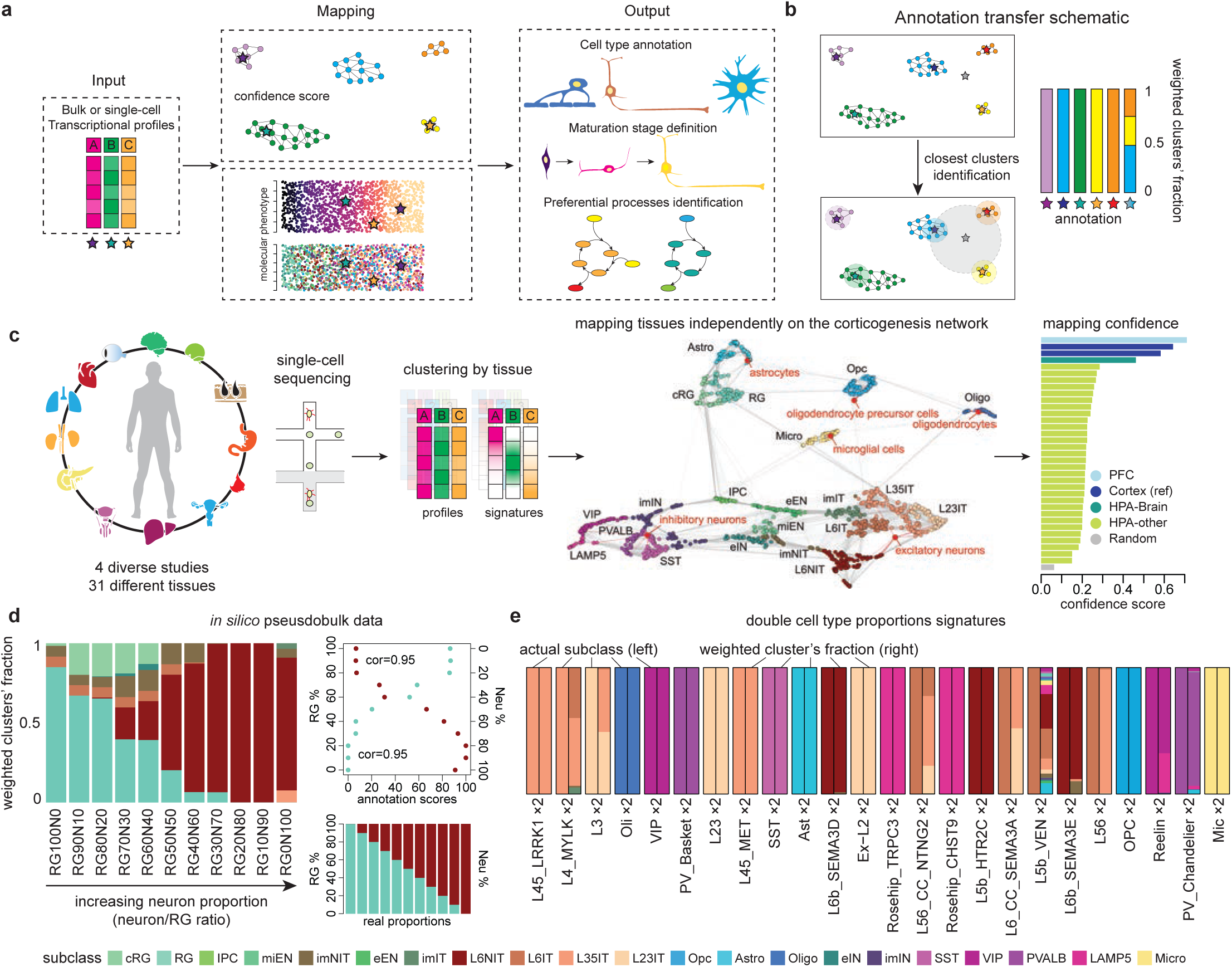
External and synthetic data validate mapping and scoring procedures. a-b. Schematics illustrating the mapping (**a**) and annotation transfer (**b**) procedures. **c** Schematic illustrating the confidence score evaluation scheme (left). Network illustrating the mapping of corresponding cortical cell types (red dots, middle). Confidence scores of all evaluation datasets mapped to the corticogenesis network colored by data source (right). **d** Annotation scores of pseudobulk profiles generated by combining increasing proportions of neurons over radial glia cells (*Methods*) (left). Relationship between annotation scores and cell composition proportions shown for radial glia (green) and neuronal (brown) cells (top plot). Data show proportions in y-axis and annotation scores in x. Composition of generated evaluation data (bottom barplot). **e** Expected subclasses (left bar) compared to annotation scores (right bar) of pseudobulk signatures in which the proportion of each annotated cell type was independently doubled (*Methods*).

## Notes

https://doi.org/10.6084/m9.figshare.30885428

### Competing Interest Statement

The authors have declared no competing interest.

